# Single-cell variant landscape of the human mitochondrial genome reveals cell type-specific functions and disease mechanisms

**DOI:** 10.64898/2026.10.11.758340

**Authors:** Chunjie Liu, Ting Peng, Zimu Cen, Peiran Lu, Prasanth Potluri, Po Hu, Shiping Zhang, Marie T. Lott, Gabrielle Widjaja, Douglas C. Wallace, Yi Xing, Liming Pei

## Abstract

Mitochondrial DNA (mtDNA) heteroplasmy is a significant source of cellular genetic diversity and can cause disease yet its distribution and functional importance across different human cell types are poorly understood. Here we present scMOCHA (Single-Cell Mitochondrial Omics for Cellular Heteroplasmy Analysis), a scalable computational framework that resolves high-confidence, single-cell resolution mtDNA heteroplasmy directly from standard single-cell RNA sequencing (scRNA-seq) data. Using this tool to interrogate publicly available scRNA-seq datasets from human peripheral blood mononuclear cells (PBMC) and bone marrow (BM) uncovered the single-cell landscape of mtDNA single-nucleotide variants (SNVs) and heteroplasmy across ages in human hematopoietic cell types. PBMCs and BM share many inherited homoplasmic and highly heteroplasmic SNVs, whereas somatic SNVs are tissue- and cell type-specific and distributed across the mtDNA genome, suggesting that they arise largely stochastically. mtDNA replication error is a dominant force shaping homoplasmic variants, while oxidative stress potentially drives a significant portion of heteroplasmic and somatic mutations. Importantly, we discover previously unknown cell type-specific and human disease-associated mtDNA SNVs. By coupling mtDNA genotypes with matched single-cell nuclear transcriptomes and structural modeling, we uncover potential molecular mechanisms by which disease-associated mtDNA SNVs impact cellular processes such as inflammation and contribute to disease. Together, these findings define the single-cell variant landscape of the human mitochondrial genome and reveal previously unrecognized lineage- and disease-associated patterns of mtDNA variation in the human hematopoietic system.

## Introduction

Mitochondria are central to cellular metabolism, bioenergetics, signaling, stress responses and organismal health. Sustained by an autonomous genome encoding core elements of the respiratory machinery^1,2^, human mtDNA exists in hundreds to thousands of copies per cell. A defining feature of mitochondrial genetics is heteroplasmy, the intracellular coexistence of mutant and wild-type (WT) mtDNA molecules^3,4^. Because heteroplasmy levels vary substantially across different cells, tissues, and developmental stages, mtDNA variation represents an important source of cellular genetic heterogeneity and a key determinant of disease penetrance. Beyond pathogenic mutations underlying inherited disorders^5^, somatic mtDNA variants accumulate progressively throughout life and have been implicated in aging and many human diseases such as neurodegeneration, inflammation, cardiovascular disease and cancer^6–11^. Emerging evidence further suggests that even low-frequency heteroplasmic variants may influence cellular fitness, metabolic states, and lineage behavior, raising the possibility that mitochondrial genetic variation may contribute to cell type-specific functional diversity across human tissues. However, a fundamental question remains unanswered: how is mitochondrial genetic heterogeneity organized across human cell types, and how is it linked to functional cellular states?

Progress in answering this question has been fundamentally bottlenecked by the inability to resolve mtDNA heteroplasmy at scale together with its single-cell transcriptional context. Genome sequence analyses revealed population-level mitochondrial genome heterogeneity^12,13^ but lack insights into mtDNA heteroplasmy at single-cell or cell-type resolution and importantly, understanding of the biological impact of mtDNA variants. Typical scRNA-seq data contain 5-25% of reads that map to mtDNA, but these mtDNA mapped reads are often filtered out or ignored during downstream analyses. Joint single-cell mtDNA and nuclear omics analyses are able to resolve mtDNA heteroplasmy at single-cell level^14,15^. However, specially modified protocols are required, and only small number of these datasets currently exist. Therefore, we currently lack robust mtDNA analysis tools at scale to utilize the rich mtDNA information often neglected in the ever-increasing, widely-used scRNA-seq datasets. Addressing this significant knowledge gap will enable understanding the broader role of mitochondrial genetic variation in shaping cellular identity, tissue homeostasis, and human disease progression.

To bridge this gap, we have developed Single-Cell Mitochondrial Omics for Cellular Heteroplasmy Analysis (scMOCHA), a scalable computational framework for integrated single-cell mitochondrial heteroplasmy and transcriptomic analysis. We use scMOCHA to interrogate approximately 5.5 million cells from all publicly available human PBMC and BM 10X scRNA-seq datasets in the NCBI GEO database (782 samples as of Nov 2025), revealing the single-cell mtDNA variant landscape of the human hematopoiesis system. Importantly, by leveraging the fact that somatic mutations occur only in certain cell lineages, we discover somatic mtDNA variants without requiring matched parental genome sequences. We find previously unknown somatic mtDNA SNVs and that somatic SNVs occur largely stochastically. Our analyses also uncover extensive lineage-specific organization of mitochondrial heteroplasmy, revealing many somatic variants enriched within distinct cellular populations and tissue compartments. By coupling mitochondrial genotypes with matched single-cell transcriptomes and through structural modeling, we further demonstrate that COVID-19-associated 8362T>G and multiple myeloma (MM)-associated 11736T>C variants impact cellular inflammatory and metabolic responses, respectively. Together, our findings reveal a previously unrecognized landscape of lineage-specific mitochondrial heterogeneity across human hematopoietic tissues, and reveal how lineage- and disease-associated mtDNA variants might impact cellular function and contribute to human disease.

## Results

### scMOCHA enables rigorous single-cell mtDNA variant detection from scRNA-seq data

To systematically identify and characterize mtDNA variants at single-cell resolution, we developed scMOCHA, a highly scalable computational framework designed for high-throughput mtDNA variant discovery and downstream analysis directly from raw 10X Genomics scRNA-seq FASTQ files, all within a single Workflow Description Language (WDL) execution (Fig. S1). Starting from raw sequencing reads, scMOCHA couples nuclear transcriptomic processing, comprising Cell Ranger alignment and automated cell-type annotation via Azimuth^16^, with a new, mtDNA-focused dual-resolution analytical module. By aggregating sparse mitochondrial reads at both individual-cell and cluster levels, this architecture unlocks high sensitivity and scalability, enabling robust variant calling by using mgatk^15^ across large single-cell cohorts. To minimize technical artifacts from sequencing errors or strand-biased alignments, the framework incorporates stringent depth and strand-concordance filtering. Integrated with MITOMAP variant annotation^17^, gnomAD^18^, and HaploGrep3 haplogroup inference^19^, scMOCHA streamlines variant annotation and automates downstream visualization including Uniform Manifold Approximation and Projection (UMAP)-based variant mapping and circular genome profiling, offering an efficient, population-scale solution for dissecting cellular mtDNA heteroplasmy.

To validate scMOCHA’s fidelity in resolving single-cell mtDNA heteroplasmy, we generated matched short-read 10X scRNA-seq (cells from 4 cell lines mixed together) and high-confidence PacBio long-read DNA-seq (each cell line sequenced separately) datasets using four human cell lines (143B, HEK293, A549, and WAL2A). We also performed the same 10X scRNA-seq in 143B, HEK293, A549, and WAL2A cell lines that lack mtDNA (Rho0 cells) to evaluate the impact of nuclear mitochondrial DNA segments (NUMTs) on the accuracy of scMOCHA.

Following quality control (QC), scMOCHA retained the majority of captured cells across both WT and Rho0 cells (Fig. S2a-c). As expected, scMOCHA detected almost no mtDNA transcript in Rho0 cells, demonstrating its ability to effectively filter NUMTs (Fig. S2b). Transcriptomic clustering clearly segregated the four distinct cell lines, demonstrating the preservation of baseline cellular identities and establishing a robust framework for benchmarking (Fig. S2d,e). Cross-platform comparison demonstrated the accuracy of scMOCHA in detecting high-confidence mtDNA SNVs from scRNA-seq data of mixed cell types (Fig. S2f). For example, in 143B cells, scMOCHA detected the same 48 SNVs from scRNA-seq as the PacBio DNA-seq. Of note, the current version of scMOCHA excludes indels, known mtDNA A-to-I RNA-editing sites, and known mis-alignment errors and homopolymer regions^20,21^ of the revised Cambridge Reference Sequence (rCRS)^22^ to ensure accuracy. In addition, short-read scRNA-seq datasets generally exhibit lower and more uneven mitochondrial genome coverage compared with long-read sequencing, limiting sensitivity for low-frequency variant detection at specific loci absent of active transcription (*e.g.*, D-loop). The 10 SNVs identified exclusively by PacBio in 143B cells were all attributable to these exclusion scenarios. Similar high concordance patterns were observed in A549, HEK293, and WAL2A cells. These benchmarking data validated the high fidelity and robustness of scMOCHA in resolving single-cell mtDNA heteroplasmy from mixed cell-type scRNA-seq data.

### The single-cell variant landscape of the mitochondrial genome in the human hematopoietic system

To systematically interrogate mitochondrial genetic heterogeneity across the human hematopoietic system, we compiled and applied scMOCHA to analyze an extensive cohort of all publicly available (as of Nov 2025) human PMBC (571 samples) and BM (211 sample) 10X scRNA-seq datasets, totaling ∼5.5 million cells after QC (Fig. 1a). These datasets are from donors of both sexes and different ages, races and ethnicities, and include both healthy and those with diseases such as COVID-19, Alzheimer’s disease (AD), MM, *etc*. (Fig. S3-4; Supplementary Table 1). Reflecting the inherent technical and biological cross-study heterogeneity of public repositories, these datasets exhibited pronounced variations in sequencing depth, mitochondrial read coverage, and donor age. Nevertheless, scMOCHA robustly resolved all major hematopoietic cell types, consistently capturing B cells, T cell subsets (CD4 T, CD8 T and other T), NK cells, monocytes (Mono), dendritic cells (DC), and hematopoietic stem and progenitor cell (HSPC, which only exists in BM samples).

**Figure 1.**
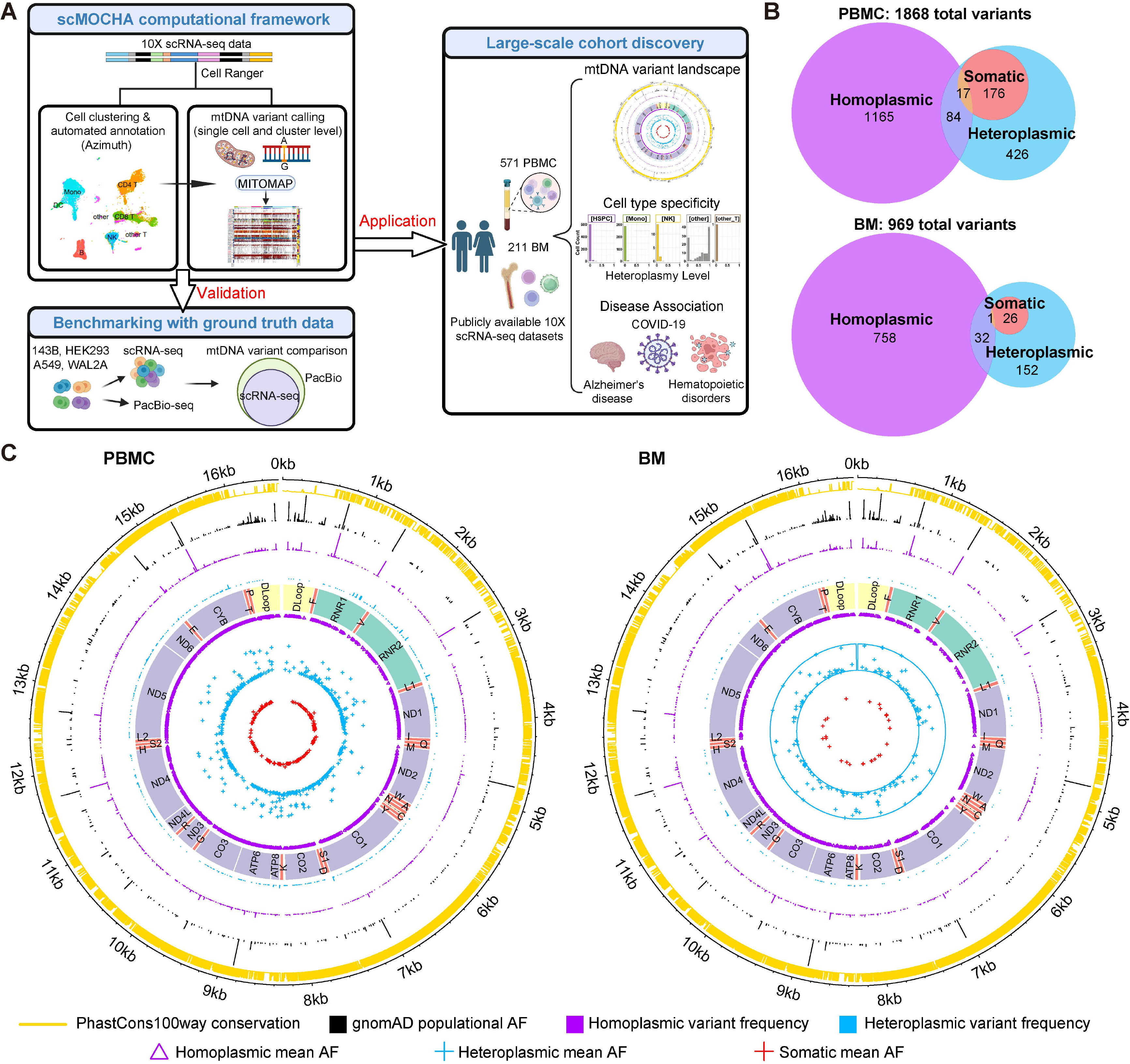
Overview of the study and mtDNA variants identified in PBMC and BM. **(a)** Overview of the study including the development and validation of scMOCHA framework and large-scale discovery workflow for single-cell mtDNA variant analysis in human PBMC and BM samples. **(b)** Venn diagrams summarize the homoplasmic (purple), heteroplasmic (blue), and somatic (red) variants identified across PBMC and BM datasets. **(c)** Circos plots illustrate the genomic distribution and allele frequency (AF) of mtDNA variants across the mitochondrial genome in PBMC and BM. Tracks from outer to inner rings are: mitochondrial genome position (black), PhastCons100way conservation (yellow), population AF derived from gnomAD (black), sample frequency of homoplasmic variants (purple) and heteroplasmic variants (blue), the mitochondrial gene annotation map (middle), mean pseudobulk AF of homoplasmic variants (purple), heteroplasmic variants (blue), and somatic variants (red).

To better decode the topology of mitochondrial genetic architecture across hematopoietic lineages, we established a hierarchical classification framework tailored for scMOCHA-derived mtDNA SNVs (Fig. S5). Following variant calling, we implemented rigorous multi-layered filtering to eliminate low-confidence candidates (fewer than 10 cells with allele frequency (AF)≥0.05 and read depth≥10) and applied aforementioned exclusion criteria. This stringent curation pipeline purged the technical artifacts typical of sparse single-cell transcriptomic reads, yielding a high-confidence mtDNA SNVs catalog of 3 categories: (1) homoplasmic (mean pseudobulk AF≥0.95 in the individual, to be consistent with definition based on genome sequencing analysis), (2) heteroplasmic (variants that are not homoplasmic, typically with 0.05≤AF<0.95), and (3) somatic, a subset of cell type-specific heteroplasmic variants (Fig. S5). Importantly, we define somatic variants as SNVs that are detected with high-confidence only in some but not all cell types for a specific donor, which indicate they are not inherited via germline. Thus, scMOCHA analyses of scRNA-seq data enabled us to identify real *de novo* somatic SNVs that would have typically required parental DNA genome analyses.

Across the 571 PBMC samples, we identified a total of 1,868 mtDNA SNVs including 193 somatic ones (Fig. 1b; Supplementary Table 2,3). In comparison, 969 total SNVs including 27 somatic variants were identified in 211 BM samples. Representative examples of homoplasmic (15734G>A), heteroplasmic (8364A>G), and somatic (7757G>A) variants at single-cell resolution demonstrated the distinct allele frequency distributions, sequencing depth profiles, and cell-type compositions characteristic of each variant class (Fig. S6). Of note, some SNVs may appear homoplasmic in some donors and heteroplasmic or somatic in other donors. 9380G>A is such an example as a homoplasmic variant in multiple donors of H1c and H6a haplogroups but a somatic variant in a donor of the J1c haplogroup (Fig. S7), underscoring the importance of haplogroup-aware interpretation of mtDNA variants.

Genome-wide distributions of these variants are shown in Circos plots (Fig. 1c). Homoplasmic variants with high AF (≥95%) are often haplogroup-associated and maternally inherited. Their distribution and frequency (purple) in both PBMC and BM match almost exactly with those from gnomAD results (black), further validating the accuracy of our analyses and general population representativeness of the samples in this study. Heteroplasmic (blue) and somatic (red) variants are distributed largely stochastically across the entire mitochondrial genome. Notably, few somatic variants were found in the D-loop, which could result from low scRNA-seq data coverage in this region due to inactive transcription. Consistent with the notion that mtDNA mutations accumulate with aging, we observed a modest yet significant positive correlation between the number of somatic variants in PBMC with donor age (Fig. S8). This was not seen in BM samples, probably due to the smaller number of somatic variants and thus lack of statistical power (Fig. S9). No significant differences in variant counts were observed between sexes except for smaller number of homoplasmic variants in male BM donors. A comprehensive heatmap visualization of all PBMC and BM variants illustrated the landscape of mtDNA, with annotations for cell type, sex, age, and mitochondrial haplogroup (Fig. S10-13).

### PBMC and BM show both shared and distinct features of mtDNA variants

Both PBMC and BM cells are of hematopoietic lineage. Comparing their mtDNA variant landscape reveals both general principles of mtDNA variation and compartment-specific features. A large portion of homoplasmic variants are shared between PBMC and BM (505 out of 1,266 PBMC and 791 BM homoplasmic variants, respectively, Fig. 2a) and their AF highly correlate with gnomAD population-level AF (Fig. S14a,b), again as they are inherited and often characteristic of specific mtDNA haplogroups. In contrast, heteroplasmic variants showed markedly less overlap (66 out of 703 PBMC and 211 BM heteroplasmic variants, respectively, Fig. 2a) and no correlation with gnomAD frequencies (Fig. S14c,d), suggesting that heteroplasmic variants are tissue-specific, likely arising through recent germline events or somatic mutagenesis in early lineages. Somatic variants were almost entirely tissue-specific, with only 3 of 193 PBMC and 27 BM somatic variants shared (Fig. 2a-c). Together with their largely universal distribution in the mitochondrial genome, this suggests that somatic mtDNA mutagenesis is tissue- and cell type-specific and largely stochastic.

**Figure 2.**
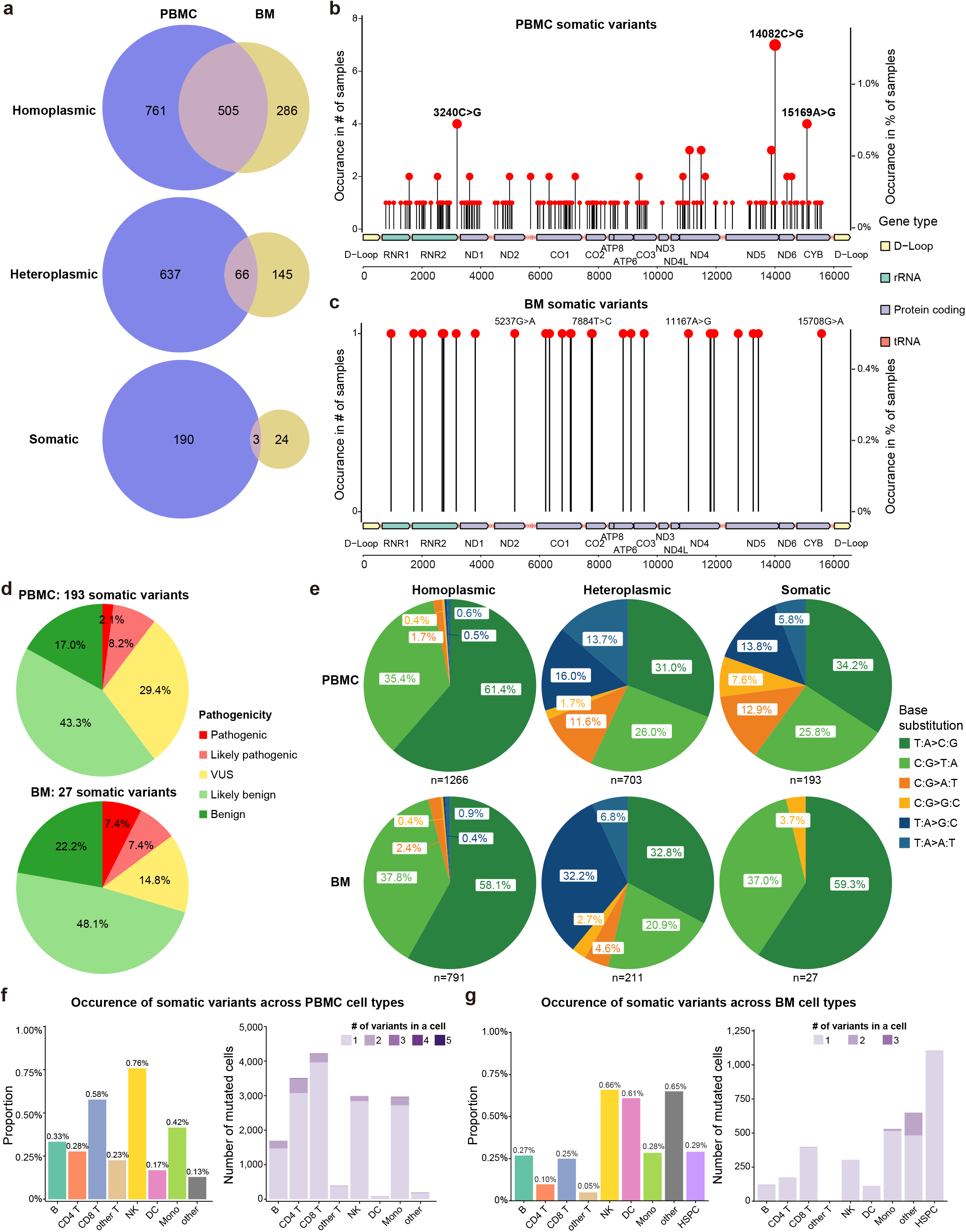
mtDNA variant landscapes and comparison between PBMC and BM. **(a)** Venn diagrams illustrate the comparison of homoplasmic, heteroplasmic, and somatic mtDNA variants identified between PBMC and BM. **(b)** Genomic distribution and AF of somatic mtDNA variants identified in PBMC. The first y-axis indicates the number of samples in which a given variant was detected, while the secondary y-axis represents the percentage of total samples harboring the variant. Annotated labels highlight recurrent hotspot variants observed across multiple unrelated individuals. **(c)** Genomic distribution and AF of somatic mtDNA variants identified in BM. The first and secondary y-axis are similar to (b). **(d)** Pathogenicity classification based on Apogee2 and nApogee of somatic mtDNA variants identified in PBMC and BM. Categories include pathogenic (red), likely pathogenic (light red), variants of uncertain significance (VUS, yellow), likely benign (light green), and benign (green). **(e)** The distribution of base substitution classes of homoplasmic, heteroplasmic, and somatic mtDNA variants identified in PBMC and BM. Colors represent base substitution classes as indicated in the legend. (**f-g**) Bar plots show the occurrence of somatic mtDNA variants within each cell type of PBMC (f) and BM (g). Left: the proportion of cells carrying somatic variants within each cell type, with percentages indicated above the bars. Right: the distribution of the number of somatic variants detected per variant-carrying cell across each cell type.

Among the 193 somatic variants identified in PBMC, several recurrent variants were detected across multiple unrelated donors. 14082C>G is the most frequent, present in 7 out of 571 or 1.2% of total samples, followed by 3240C>G and 15169A>G that occur in 4 out of 571 or 0.7% of total samples (Fig. 2b). Considering the largely random nature of somatic mutagenesis and that these donors belong to different mtDNA haplogroups and are therefore genetically unrelated (Fig. S15), this high percentage recurrence is unexpected and suggests existence of potential somatic mutation hotspots. With much fewer available BM samples and somatic variants, we did not observe similar somatic mutation hotspots in BM (Fig. 2c).

Pathogenicity assessment using the APOGEE2^23^ and nAPOGEE^24^ prediction tools revealed that the majority of PBMC and BM somatic variants were predicted as benign or likely benign (Fig. 2d; Supplementary Table 4). However, 10.3% PBMC and 14.8% BM somatic variants are predicted to be pathogenic or likely pathogenic. These include known, rare pathogenic mutation 3890G>A (*MT-ND1* p.R195Q) which causes a spectrum of Leber hereditary optic neuropathy (LHON) and Leigh-like encephalopathy^25,26^, and likely pathogenic variant 13376T>C (*MT-ND5* p.I347T) which was reported to be associated with mitochondrial encephalomyopathy, lactic acidosis, and stroke like episodes (MELAS)^27^, supporting confidence of the predictions. Overall, these results suggest that although a relatively small fraction, pathogenic and likely pathogenic somatic mutations do arise in the human mitochondrial genome in multiple tissues to detectable levels.

Recent genome DNA-seq analyses suggested that mtDNA replication errors rather than oxidative stress likely drive age-related mtDNA mutation accumulation^28–30^, based on the base substitutions being transitions (most likely caused by DNA replication errors) or transversions (most likely caused by oxidative stress). In both PBMC and BM, we found that germline homoplasmic variants were dominated by T:A>C:G and C:G>T:A transitions (>96%) (Fig. 2e; Fig. S16), consistent with the well-characterized strand-asymmetric mutational bias resulting from mtDNA replication errors and confirming these recent discoveries^28–30^. In contrast, heteroplasmic and somatic variants exhibited a more diverse base substitution spectrum. Transversion base substitutions reach over 40% for heteroplasmic variants in both PBMC and BM, and for somatic variants in PBMC. These results suggest that although mtDNA replication error is most likely the dominant force shaping homoplasmic variants, oxidative stress may drive a significant portion of heteroplasmic and somatic mutations in the mtDNA genome.

Examination of somatic variant distribution across PBMC and BM cell types revealed differential cell-type enrichment (Fig. 2f,g; Supplementary Table 5). Overall, less than 1% of cells carry somatic variants across all cell types. In PBMC, several lymphocyte cell types exhibited high proportion of cells carrying somatic variants, with the highest in NK cells (0.76%), followed by CD8 T cells (0.58%), monocytes (0.42%), B cells (0.33%), and CD4 T cells (0.28%). In BM, NK cells exhibited the highest proportion of somatic variants (0.66%), followed by other cells (0.65%) and DCs (0.61%). Most somatic variant-carrying cells have a single somatic variant, though a small percentage of cells harbor multiple and up to five co-occurring somatic variants.

Per-locus enrichment analysis of non-haplogroup variants between PBMC and BM revealed significant tissue-specific differences in the genomic distribution of heteroplasmic variants (Fig. S17). *MT-RNR1* and *MT-RNR2* (rRNA genes) were significantly enriched for heteroplasmic variants in PBMC relative to BM, while *MT-ATP6*, *MT-ND3*, *MT-ND4L*, and *MT-ND4* showed significant enrichment of heteroplasmic variants in BM. No significant tissue-specific enrichment was observed for homoplasmic or somatic variant categories.

### mtDNA variants with cell type-specific heteroplasmy enrichment and their transcriptional consequences

A few recent studies have identified potentially cell type-specific purifying selection of mtDNA variants. For example, 3243A>G associated with MELAS shows broad range of heteroplasmy across most PBMC cell types but markedly reduced heteroplasmy in T cells in some MELAS patients^31^. The synonymous variant 7076A>G (*MT-CO1*, p.G391=) exhibited CD8 effector memory T cell-specific selection against the 7076G heteroplasmy but not in other T cell subtypes in a healthy donor^32^. We examined such cell type-selective high or low mtDNA variant heteroplasmy enrichment phenomenon and identified a comprehensive catalog of mtDNA variants with major cell type-specific heteroplasmy enrichment in both PBMC and BM (Fig. S18-19; Supplementary Table 6). These variants spanned the entire mitochondrial genome, affecting rRNA, tRNA and protein-coding genes, and displayed diverse pathogenicity predictions ranging from benign to likely pathogenic. Many variants exhibited striking cell type specificity, with high mean AF and cell proportions restricted or depleted in specific cell types.

For example, 10500G>A, a rare variant in population^17^, was identified as a heteroplasmic variant in PBMC samples with high AF predominantly found in B cells and NK cells, and all 3 T cell subtypes (CD4, CD8, and other T) showed very low heteroplasmy (Fig. S20a). This variant results in a nonsynonymous substitution from alanine to threonine at position 11 of the ND4L protein (*MT-ND4L*, p.A11T), a subunit of mitochondrial Complex I. The scRNA-seq data enabled us to investigate the biological impact of this variant on nuclear gene expression by comparing high (AF≥0.5) *vs* low (AF<0.5) heteroplasmy cells of the same cell type from the same donor. This strategy minimizes donor-specific genetic and cell-type effects and focuses on the transcriptional differences primarily driven by 10500G>A heteroplasmy. B cells with high heteroplasmy (AF≥0.5) 10500G>A exhibited upregulation of genes enriched in response to xenobiotic stimulus, B cell receptor signaling, Fc-epsilon receptor signaling, immune response-regulating cell surface receptor signaling, antigen receptor-mediated signaling, and lymphocyte activation (Fig. S20b, left), and downregulation of genes in antimicrobial humoral response, antibacterial humoral response, complement activation (classical pathway), adaptive immune response, defense response to bacterium, and plasma membrane organization (Fig. S20b, right). In NK cells, high 10500G>A heteroplasmy was associated with upregulation of T cell activation, leukocyte cell-cell adhesion, regulation of cell adhesion, lymphocyte activation, and adaptive immune response based on somatic recombination (Fig. S20c, left), while downregulated genes were enriched in lymphocyte mediated immunity, immune effector process, cell killing, leukocyte mediated cytotoxicity, and positive regulation of programmed cell death (Fig. S20c, right). These results demonstrate that 10500G>A heteroplasmy levels modulate distinct immune effector programs in a cell type-specific manner – promoting receptor signaling while dampening humoral responses in B cells, and enhancing cell adhesion and activation while suppressing cytotoxic function in NK cells.

14831G>A is another example that may modulate cell type-specific immune function. It showed preferential high heteroplasmy only in CD4 and CD8 T cells in one donor PBMC sample (Fig. S21a). This variant results in a nonsynonymous substitution from alanine to threonine at position 29 (A29T) of cytochrome b (*MT-CYB*, p.A29T), a core subunit of mitochondrial Complex III. In CD4 T cells with high heteroplasmy (AF≥0.5), upregulated genes were enriched in cell killing, immune effector process, leukocyte mediated cytotoxicity, leukocyte mediated immunity, adaptive immune response, phospholipase C-activating GPCR signaling, inflammatory response, and cell-cell adhesion (Fig. S21b, left). Downregulated genes were prominently enriched in cytoplasmic translation, mononuclear cell differentiation, lymphocyte differentiation, T cell differentiation, ribosomal small subunit biogenesis, ribosome biogenesis, T cell activation, and T cell receptor signaling (Fig. S21b, right). In CD8 T cells, a similar but distinct pattern emerged: upregulated genes were enriched in disruption of cells of another organism, cell killing, cellular defense response, positive regulation of cytosolic calcium ion concentration, and chemokine-mediated signaling (Fig. S21c, left), while downregulated genes were associated with adaptive immune response, lymphocyte activation, T cell activation, lymphocyte differentiation, and regulation of T cell activation (Fig. S21c, right). Together, these findings indicate that 14831G>A heteroplasmy in *MT-CYB* promotes a transcriptional shift toward cytotoxic effector programs while suppressing T cell differentiation and translational machinery in both CD4 and CD8 T cell compartments, consistent with a terminal effector phenotype potentially driven by Complex III-mediated metabolic perturbation.

### 8362T>G heteroplasmy is decreased in COVID-19 and associated with monocyte inflammatory reprogramming

Many mtDNA variants are known to be pathogenic as the driver mutation of specific mitochondrial disease. However, whether and how certain mtDNA variants can serve the “passenger” role to modify disease susceptibility or response remain much less understood. Comparing specific variant occurrence and heteroplasmy levels in various diseases *vs* healthy donors of the same dataset, we identified multiple variants as potential disease modifiers.

A prominent example is 8362T>G, located in the *MT-TK* gene encoding mitochondrial tRNA-Lysine. This variant is rare in human population^17^ but detected as a heteroplasmic variant across multiple PBMC samples from a COVID-19 dataset and exhibited a striking disease-specific distribution – all cell types in the COVID-19 patients showed significantly reduced heteroplasmy levels than controls (Fig. 3a).

**Figure 3.**
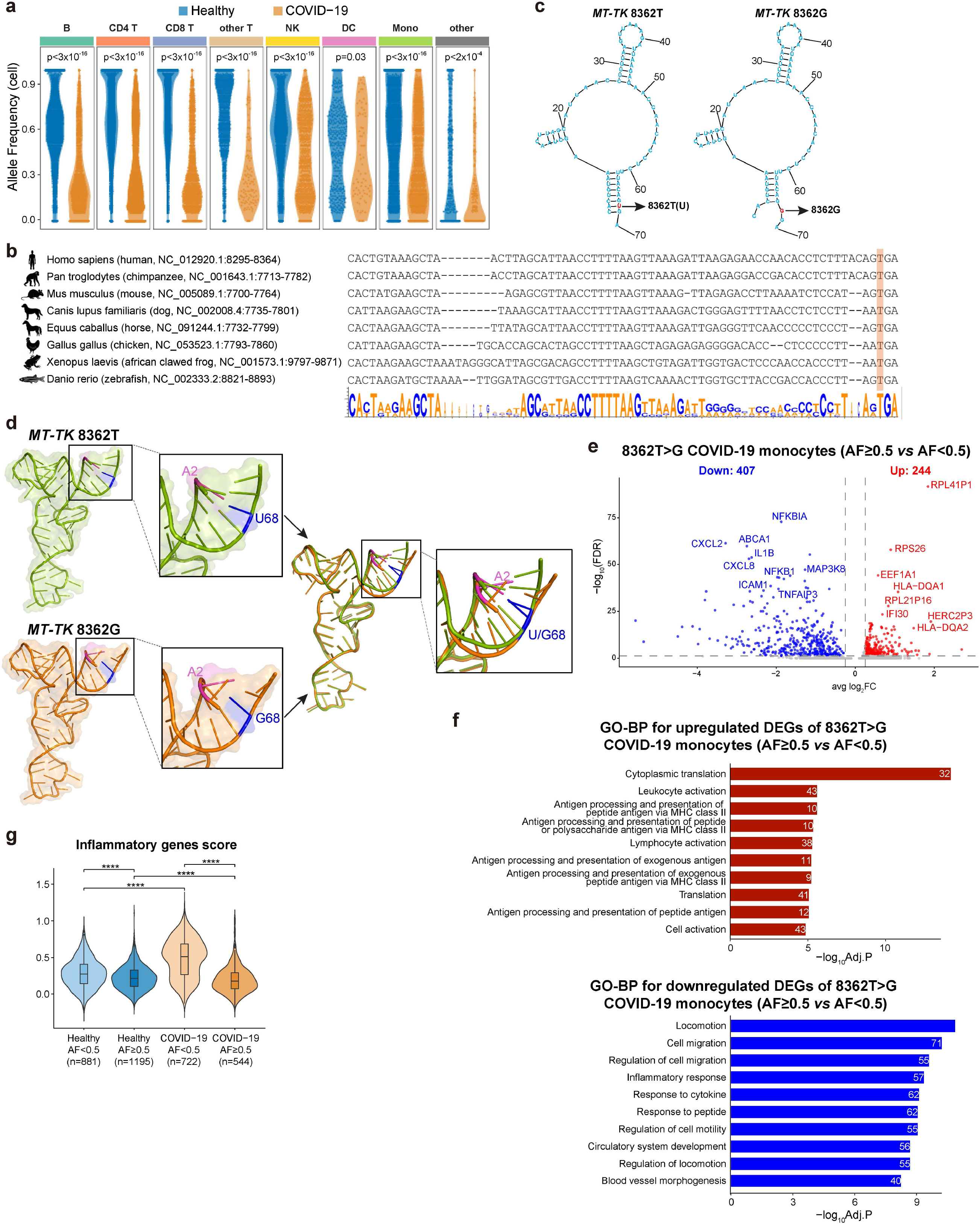
Monocyte inflammatory reprogramming associated with 8362T>G (*MT-TK*) in COVID-19. **(a)** Violin plots comparing the single-cell allele frequency (AF) distribution of the 8362T>G variant between healthy controls (blue) and COVID-19 patients (orange) across major PBMC cell types, including B cells, CD4 T cells, CD8 T cells, other T cells, NK cells, dendritic cells (DC), monocytes (Mono), and “other” cell types. P-values from inter-group comparisons are shown above each panel. **(b)** Evolutionary conservation analysis of human *MT-TK* 8362T across representative vertebrate and invertebrate species. The sequence logo below summarizes nucleotide conservation across aligned sequences, demonstrating strong evolutionary constraint within the acceptor stem region of *MT-TK*. **(c)** Predicted RNA secondary structures of the WT *MT-TK* tRNA (8362T, left) and the mutant tRNA (8362G, right). The 8362T>G substitution occurs at position 68 within the acceptor stem of the tRNA-Lysine molecule. **(d)** Three-dimensional structural models of the WT *MT-TK* 8362T (top) and mutant *MT-TK* 8362G (bottom) generated by RNAComposer. Zoomed-in views highlight the canonical A2-U68 Watson-Crick base pair in the WT structure and the A2-G68 mismatch introduced by the T>G substitution, which disrupts the local geometry of the acceptor stem at the critical 5ʹ-3ʹ junction. **(e)** Volcano plot showing differentially expressed genes (DEGs) in COVID-19 monocytes with high 8362T>G heteroplasmy (AF≥0.5) compared to those with low heteroplasmy (AF<0.5). Significantly upregulated genes (n=244) are shown in red and downregulated genes (n=407) in blue. Dashed lines indicate thresholds for statistical significance (adjusted P<0.05) and fold change (|avg log₂FC|>0.25). **(f)** Gene Ontology Biological Process (GO-BP) enrichment analysis of upregulated (top) and downregulated (bottom) DEGs in high-heteroplasmy COVID-19 monocytes. Bar length represents enrichment significance as −log_10_(adjusted P-value), and numbers inside each bar indicate the number of genes associated with the corresponding biological process. **(g)** Gene set scoring analysis of inflammatory gene expression across four monocyte populations stratified by disease status and 8362T>G heteroplasmy level: healthy with low AF (AF<0.5), healthy with high AF (AF≥0.5), COVID-19 with low AF (AF<0.5), and COVID-19 with high AF (AF≥0.5). Cell numbers for each group are indicated. **** P < 0.0001.

8362T is highly conserved across species from zebrafish to human (Fig. 3b). The 8362T>G substitution occurs at position 68 of the *MT-TK* gene within the acceptor stem of the tRNA secondary structure, changing the WT uracil (U68) to guanine (G68) (Fig. 3c). In the WT tRNA, U68 forms a canonical Watson-Crick A2-U68 base pair that stabilizes the acceptor stem. Structural modeling suggest that the T>G substitution disrupts this pairing and alters the local geometry of the acceptor stem, with the A2-G68 mismatch introducing structural distortion at the critical 5ʹ-3ʹ junction of the tRNA (Fig. 3d). Such perturbations in the acceptor stem are known to affect tRNA aminoacylation efficiency and stability, potentially leading to mitochondrial translational stress.

To further investigate the potential functional consequences of 8362T>G variant, we performed differential gene expression analysis comparing COVID-19 monocytes with high heteroplasmy (AF≥0.5) to those with low heteroplasmy (AF<0.5). This comparison identified 244 significantly upregulated and 407 significantly downregulated genes (Fig. 3e). Gene Ontology biological process (GO-BP) enrichment analysis revealed that upregulated genes in high-heteroplasmy monocytes were predominantly enriched in cytoplasmic translation, leukocyte activation, and multiple antigen processing and presentation pathways (Fig. 3f, top), such as *RPL41P1*, *RPS26*, *EEF1A1*, *HLA-DQA1*, *RPL21P16*, *IFI30*, *HERC2P3*, and *HLA-DQA2* (Fig. 3e). Notably, previous studies have suggested that disturbance of mitochondrial translation alters cytoplasmic translation^33^.

Conversely, downregulated genes in high-heteroplasmy monocytes were enriched in pathways associated with locomotion, cell migration, regulation of cell migration, inflammatory response, response to cytokine, response to peptide, and regulation of cell motility (Fig. 3f, bottom), indicating a suppression of classical inflammatory and migratory programs. These include many key NF-κB signaling components and inflammatory mediators – *NFKBIA*, *NFKB1*, *IL1B*, *CXCL8*, *CXCL2*, *MAP3K8*, *ICAM1*, and *TNFAIP3* (Fig. 3e). These transcriptional changes were further corroborated by gene set scoring analysis across four cell populations stratified by disease status and heteroplasmy level (Fig. 3g). COVID-19 patient monocytes with low 8362T>G heteroplasmy (AF<0.5) showed significantly increased inflammatory gene scores (based on 30 curated inflammatory genes^34^) compared to healthy donor monocytes with low 8362T>G heteroplasmy, likely reflecting COVID-19-induced inflammatory response. However, presence of high 8362T>G heteroplasmy (AF≥0.5) in COVID-19 patient monocytes significantly dampened such inflammatory response, suggesting 8362T>G heteroplasmy may act as a genetic modifier and is associated with monocyte inflammation reprogramming in COVID-19.

### 11736T>C is associated with plasma cell mitochondrial dysfunction and structural perturbation of Complex I in MM

Another human disease-associated mtDNA variant is 11736T>C. This variant results in a nonsynonymous amino acid substitution from leucine (L) to proline (P) at position 326 (L326P) in the ND4 protein, a core subunit of mitochondrial Complex I. This Leucine is highly evolutionally conserved (Fig. 4a). 11736T>C was found as a heteroplasmic variant in a BM sample of a donor newly diagnosed with MM (GEO sample: GSE271107-GSM8369878), a cancer of the plasma B cells in the BM. This variant showed high heteroplasmy predominantly in plasma B cells of the “other” cell population, while few other cell types contain significant number of high heteroplasmy cells (Fig. 4b,c).

**Figure 4.**
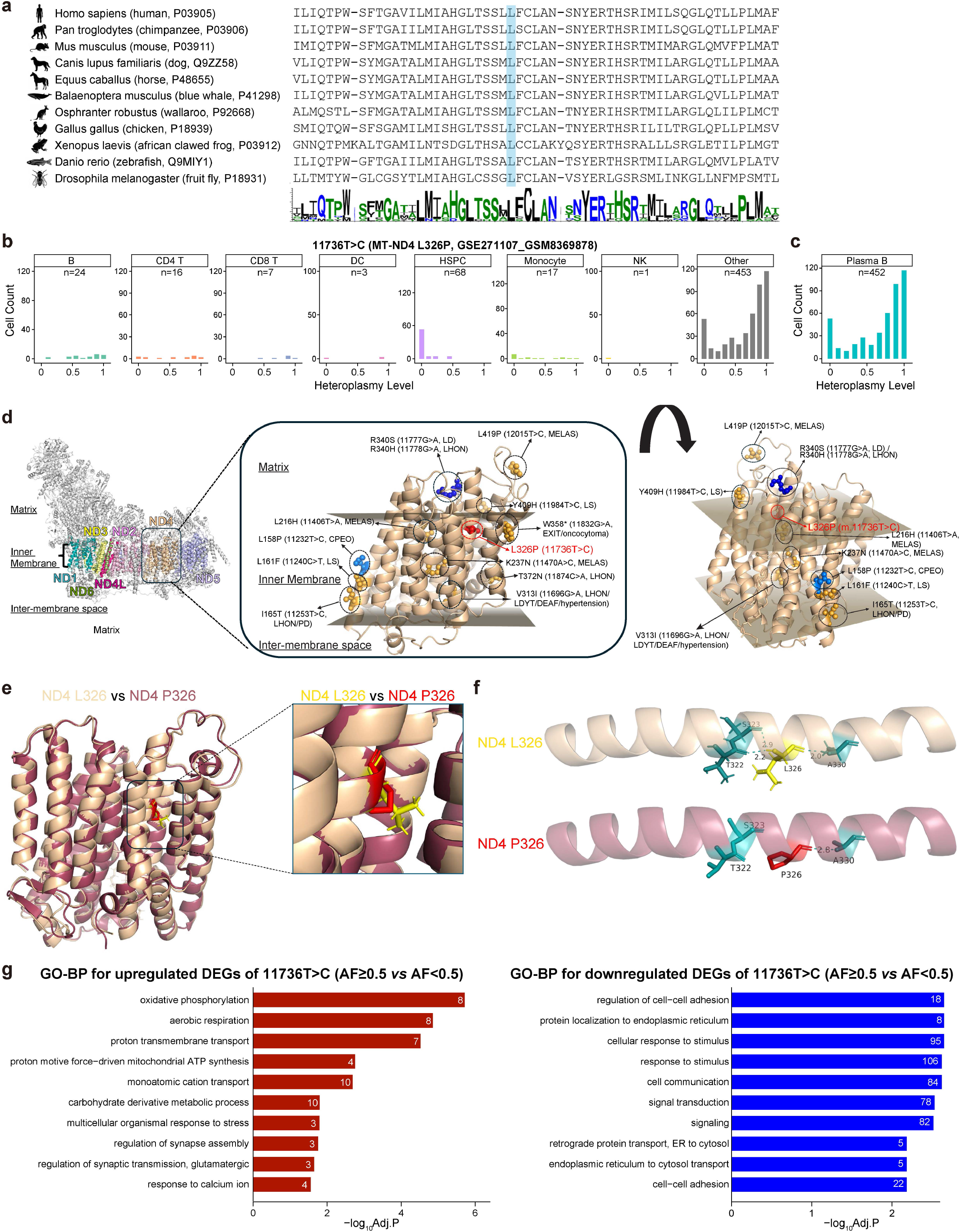
Cell type-specific enrichment and structural characterization of the 11736T>C (*MT-ND4*, p.L326P) variant in multiple myeloma. (**a**) Evolutionary conservation analysis of the ND4 L326 residue across representative vertebrate and invertebrate species. The sequence logo below summarizes residue conservation across aligned sequences, demonstrating high evolutionary constraint in this ND4 transmembrane region. (**b**) Histograms showing the allele frequency distribution of the variant 11736T>C within annotated cell types, including B cells, CD4 T cells, CD8 T cells, other T cells, dendritic cells (DC), hematopoietic stem and progenitor cells (HSPC), monocytes, NK cells, and “other” cells. **(c)** The “other” cells in this sample are almost entirely plasma B cells (452 out of 453 cells). Histograms show the allele frequency distribution of the variant 11736T>C within plasma B cells. **(d)** Structural localization of 11736T>C within the ND4 subunit of Complex I (PDB: 9I4I). The left panel shows an overview of Complex I, with different subunits labeled in distinct colors. The middle panel presents a zoomed-in side view of ND4, highlighting previously reported disease-associated residues, most shown in black, R340S/R340H in dark blue, L158P in light blue, and L326P in red. The right panel shows a corresponding top view of the ND4 zoom-in. **(e)** Structural model comparison of ND4 L326 (colored wheat) and P326 (colored raspberry). The left panel shows the overall structural alignment, while the right panel provides a zoomed-in view highlighting local geometric changes at residue 326, with L326 shown in yellow and P326 in red. **(f)** Hydrogen bond interactions of residue 326 with nearby amino acids in ND4-L326 (top) and ND4-P326 (bottom). L326 is shown in yellow, P326 in red, and surrounding residues in teal. Hydrogen bonds are indicated by dashed lines, and residues are labeled in black. Distances (Å) between residues are indicated. **(g)** Gene Ontology Biological Process (GO-BP) enrichment analysis of upregulated (left) and downregulated (right) differentially expressed genes (DEG) associated with high 11736T>C allele frequency (AF≥0.5 vs AF<0.5).

L326 is located in an inner-membrane α-helix of the ND4 protein of Complex I (Fig. 4d; PDB 9I4I), a hotspot region containing many known pathogenic mtDNA SNVs, including the well-characterized 11778G>A mutation (causing LHON; dark blue) and others associated with Leigh syndrome and MELAS, etc. Interestingly, another leucine-to-proline substitution nearby, *MT-ND4* p.L158P (11232T>C; light blue), has previously been reported to cause chronic progressive external ophthalmoplegia (CPEO)^35^ and susceptibility to severe neurologic events after general anesthesia^36^.

Proline substitutions within α-helices often result in significant structural change and are commonly referred to as “helix breakers” because the cyclic side chain of proline restricts backbone flexibility and destabilizes helical conformations. We therefore next explored the potential structural impact of ND4 L326P on mitochondrial Complex I. Structural modeling using AlphaFold3-generated P326 variant structure compared with the experimentally resolved WT L326 structure suggested that the global ND4 fold remained largely preserved, with a relatively small backbone Root-Mean-Square Deviation (RMSD) of 0.58Å between L326 and P326 structures (Fig. 4e). However, detailed local analysis revealed subtle yet significant conformational perturbations surrounding residue 326 within the transmembrane region. Specifically, the L326P substitution introduced increased local helix bending and disrupted nearby backbone hydrogen bond interactions that normally stabilize the α-helical structure (Fig. 4f). Similar structural analyses performed for the reported pathogenic L158P mutation showed comparable hydrogen-bond disruption patterns (Fig. S22a), further supporting the structural sensitivity of leucine-to-proline substitutions within ND4 transmembrane helices. Additional geometric analysis demonstrated modest but reproducible changes in helix angle in both the L326P and L158P mutant structures (Fig. S22b,c), consistent with local distortion induced by proline-mediated helix destabilization. These structural perturbations occur within the inner-membrane region of ND4 and may impact Complex I function by influencing local conformational dynamics or proton translocation efficiency.

These structural insights were further supported by gene expression changes in plasma B cells with high 11736T>C heteroplasmy (AF≥0.5) compared to those with low heteroplasmy (AF<0.5) (Supplementary Table 7). The high heteroplasmy cells showed significantly increased expression of many genes in oxidative phosphorylation and aerobic respiration pathways (Fig. 4g; Fig. S22d), indicating compensatory responses to mitochondrial stress. In addition, several genes known to be associated with MM progression and tumor biology, including *DKK1*, *CAMK2D*, and *MEF2C*, were also upregulated. Notably, DKK1 has been widely reported as a biomarker and potential therapeutic target in MM, with elevated expression associated with tumor progression and metastasis in multiple cancer types^37^. In contrast, downregulated genes in 11736T>C high heteroplasmy plasma B cells were mainly enriched in pathways associated with cell-cell adhesion and signaling, ER stress. Together, these findings identify 11736T>C (*MT-ND4*, p.L326P) as a plasma B cell-enriched mtDNA variant associated with substantial mitochondrial transcriptional remodeling and localized structural perturbation of Complex I, suggesting a potential role of mitochondrial dysfunction and metabolic adaptation in MM.

## Discussion

Although mtDNA variation has long been implicated in aging, metabolism, immunity, and human disease, most previous studies have relied on bulk sequencing approaches that average mitochondrial heterogeneity across complex cellular populations. Consequently, the organizational principles governing mtDNA variation at single-cell resolution have remained poorly understood. In this study, we develop scMOCHA, a scalable computational framework for systematic interrogation of mitochondrial genetic variation from scRNA-seq datasets and uncover the single-cell landscape of the mitochondrial genetic heterogeneity in the human hematopoietic system. Our findings reveal that mitochondrial genetic variation is highly structured across tissues and lineages and establish mitochondrial heteroplasmy as a dynamic and context-dependent component of cellular identity.

scMOCHA introduces several innovations that, in combination, distinguish it from existing mtDNA variant calling tools. First, it couples reference-aware mtDNA alignment against the rCRS with chemistry-adaptive single-cell quality control and Azimuth-based automatic reference cell-type annotation, so that variant calling is performed on cell type-stratified pseudobulks as well as on millions of single cells in the same run. Second, it implements a transparent three-step variant-classification framework that explicitly filters mis-alignment-prone positions and RNA editing sites, partitions high-confidence variants into homoplasmic and heteroplasmic strata and nominates somatic mutations only when a heteroplasmic variant displays a cell type-restricted pattern by non-parametric testing, which is substantially more rigorous than simply based on allele-frequency thresholds. Third, it integrates HaploGrep3 haplogroup inference, MITOMAP locus and disease annotation, and gnomAD population frequencies into a unified cell-by-variant table backed by a DuckDB columnar database, enabling cross-cohort queries over tens of millions of cell-by-variant observations. Fourth, it tightly couples variant calls to phenotype: cells carrying a nominated mtDNA variant can be directly stratified by heteroplasmy level for differential gene expression and pathway enrichment analyses against same cell types from the same donor, thereby linking mitochondrial genotype to lineage-specific transcriptional state in a single experiment. Our findings further demonstrate that mtDNA variant states are highly context dependent, as the same variant can exhibit homoplasmic, heteroplasmic, or somatic characteristics in distinct donor or lineage contexts. These observations challenge traditional static categorizations of mtDNA mutations and instead support a continuum model of mitochondrial genetic states shaped by cellular and tissue environments.

One of the most striking findings of our study is the hierarchical organization of mtDNA variation across hematopoietic tissues. Homoplasmic variants exhibit strong conservation between PBMC and BM compartments and correlation with gnomAD results, largely reflecting inherited, population-level mitochondrial genetic backgrounds. In contrast, heteroplasmic variants show pronounced tissue partitioning, whereas somatic variants display highly individualized and lineage-restricted distributions with minimal overlap between PBMC and BM. The broad distribution of somatic variants across the mitochondrial genome and their little overlap between PBMC and BM suggest that somatic variants arise through local lineage-specific events in a largely stochastic fashion.

Interestingly, distinct mtDNA variant classes also exhibit markedly different mutational signatures and strand asymmetries, suggesting potentially different underlying mutational mechanisms. Homoplasmic variants are predominantly characterized by classical replication-associated nucleotide transition signatures, consistent with long-term accumulation of mtDNA replication errors. In contrast, heteroplasmic variants in PBMC and BM, and somatic variants in PBMC, demonstrate increased nucleotide transversion associated with oxidative stress and reactive oxygen species (ROS)-related mitochondrial damage. Notably, somatic variants in BM samples partially revert toward replication-associated mutational signatures, suggesting that mitochondrial mutagenesis may differ substantially across tissue microenvironments and hematopoietic cellular states. These observations support the concept that mitochondrial heteroplasmy is not merely stochastic genetic noise, but may instead reflect dynamic interactions between replication fidelity, oxidative metabolism, and lineage-associated selective pressures.

Another major finding of our study is the strong lineage-associated organization of mitochondrial heterogeneity across immune cell types. Among hematopoietic cell types, NK cells consistently exhibited the highest proportions of somatic mtDNA variants in both PBMC and BM datasets. Interestingly, most mutated cells contain only a single detectable somatic mutation, suggesting that mitochondrial somatic mutagenesis in the hematopoietic system is generally sparse rather than massively accumulated. Given the highly cytotoxic and metabolically active nature of NK cells, these observations raise the possibility that mitochondrial stress, activation-associated ROS production, or rapid cellular turnover may contribute to enhanced mitochondrial genetic diversification within this lineage. More broadly, our results suggest that mitochondrial heteroplasmy is closely linked to hematopoietic cell functional states and may represent an additional layer of hematopoietic cell heterogeneity that cannot be resolved through nuclear transcriptomics alone.

Our analyses further reveal strong associations between lineage-enriched mitochondrial heteroplasmy and hematopoietic cell transcriptional and potentially functional remodeling. High heteroplasmy of certain variants such as 10500G>A and 14831G>A are enriched or depleted in specific cell types and are associated with altered cytotoxicity, cytokine signaling, leukocyte activation, and translational programs in related cell types. Although these observations do not establish direct causality, they suggest that mitochondrial genetic heterogeneity may be closely integrated with immune cell functional states. Given the central role of mitochondria in cellular energetics, redox regulation, and innate immune signaling, these findings raise the possibility that mitochondrial genetic variation may contribute to shaping immune cell phenotypic diversity in both physiological and pathological contexts.

The disease-associated mtDNA variants identified in this study further support the functional relevance of mitochondrial genetic heterogeneity in human disease. In COVID-19 samples, the *MT-TK* variant 8362T>G exhibits substantially reduced heteroplasmy levels across cell types compared with healthy controls, and high heteroplasmy is strongly correlated with reduced inflammatory response in COVID-19. We also identified the *MT-ND4* variant 11736T>C existing in high heteroplasmy only in plasma B cells from an MM donor. High heteroplasmy of this variant is associated with substantial mitochondrial transcriptional remodeling and localized structural perturbation of Complex I. Both these variants and those with cell type-specific heteroplasmy enrichment discussed above demonstrate a general biological principle: mtDNA variants are an important component of human genetic heterogeneity, and they are cell type-specific or enriched and substantially impact mitochondrial biology and specific cellular functions. The discovery of potential mtDNA somatic mutation hotspot (*e.g.*, 14082C>G in PBMC) in multiple unrelated individuals is interesting. It is tempting to compare this to clonal hematopoiesis^38,39^, the phenomenon of somatic nuclear DNA mutations in certain “hotspot” genes (*e.g.*, *DNMT3A*, *TET2*, *ASXL1*, *JAK2*) conferring a fitness or growth advantage on the mutant cell in the hematopoietic system. Clonal hematopoiesis is particularly prevalent in the elderly population. Even clonal hematopoiesis of indeterminate potential (CHIP) is associated with increased risk of various diseases such as blood cancer and cardiovascular disease. As mtDNA mutations accumulate with aging too, it would be interesting to determine whether specific mtDNA mutation itself drives clonal hematopoiesis-like phenomenon and if so, whether this underlies many aging-associated human diseases. Compared to studies of CHIP, the sample size here remains too small to address this question, even though we have investigated all currently available public PBMC and BM 10X scRNA-seq datasets. With the fast-growing number of scRNA-seq datasets, this question will be revisited in the future to reveal new biological insights into how mtDNA variation impacts cellular and organismal physiology.

Several limitations of this study warrant consideration. First, although scMOCHA incorporates extensive filtering procedures to reduce NUMT contamination and sequencing artifacts, complete elimination of NUMT interference remains challenging for short-read scRNA-seq datasets. Second, the current implementation of scMOCHA does not support mitochondrial indel detection, potentially underestimating the full spectrum of mitochondrial genetic variation. Third, some disease-associated variants were identified in small donor subsets; larger cohorts with variant curation using American College of Medical Genetics and Genomics (ACMG) standards^40^ will be required to determine their recurrence and potential pathogenic significance. Our current analyses focused primarily on PBMC and BM tissues. Future applications of scMOCHA to other organs such as heart, brain, kidney, and skeletal muscle may further reveal tissue-specific mtDNA landscapes. Finally, although our study identifies strong associations between mtDNA heteroplasmy and cellular transcriptional states, direct experimental perturbation studies will be needed to determine whether specific mitochondrial variants causally impact immune remodeling, metabolic adaptation, or disease progression.

In summary, our study establishes a single-cell framework for systematic interrogation of mitochondrial genetics across human tissues, defines the single-cell variant landscape of the human mitochondrial genome in the human hematopoietic system, and provides evidence that mitochondrial heteroplasmy represents a dynamic, lineage-associated, and biologically structured component of human genetic heterogeneity.

## Methods

### Collection of publicly available scRNA-seq datasets

Publicly available scRNA-seq datasets generated using the 10X Genomics Chromium platform with multiple library chemistries, including 3′ (SC3Pv2/v3/v3.1) and 5′ (SC5P-PE and SC5P-R2), were collected from the Gene Expression Omnibus (GEO). A total of 571 human PBMC samples and 211 BM samples, covering healthy individuals and a broad range of diseases such as Alzheimer’s disease, COVID-19, and hematopoietic disorders, were included in this study. For BM datasets, samples associated with transplantation or severe technical abnormalities were excluded to minimize potential confounding effects. Metadata, including tissue source, disease status, sequencing chemistry, donor age, sex, and sample annotations, were manually curated and summarized. Donor sex was further confirmed using the expression of sex-specific markers (*e.g.*, *XIST* and Y chromosome genes). Sankey plots were subsequently generated to visualize the relationships among metadata categories and the overall sample composition across studies.

### Overview and development of scMOCHA pipeline

A customized computational pipeline, termed scMOCHA (<u>S</u>ingle-<u>C</u>ell <u>M</u>itochondrial <u>O</u>mics for <u>C</u>ellular <u>H</u>eteroplasmy <u>A</u>nalysis), was developed in this study as an end-to-end workflow that converts raw 10X Genomics scRNA-seq reads into per-cell and per-cluster mitochondrial heteroplasmy profiles, together with automated cell-type annotation and variant pathogenicity interpretation. The pipeline was implemented using WDL and executed with Cromwell on either local workstations or Simple Linux Utility for Resource Management (SLURM)-managed high-performance computing (HPC) clusters. To ensure reproducibility across computational environments, all dependencies were version-controlled within Conda environments and additionally distributed through Docker/Singularity containers (Xinglab/scMOCHA).

### Pipeline implementation and computational resource requirements

The scMOCHA WDL pipeline was validated under Cromwell (v78). Resource configurations were managed via dedicated deployment blueprints. Individual sample was processed with a standard baseline allocation of 10 CPU cores, 50 GB of RAM, and 50 GB of scratch storage. To ensure strict environmental reproducibility, all software dependencies were encapsulated within a Conda environment configuration (env.prod.yaml) and compiled into a versioned Docker image (Xinglab/scMOCHA), which can be converted natively into a Singularity image archive (.sif) for secure HPC execution. Pipeline reentrancy and intermediate file caching to support partial task restarts were managed via an integrated HyperSQL database (HSQLDB) backend.

### Mitochondrial read alignment

Raw FASTQ files were aligned and demultiplexed using Cell Ranger (v8.0.0) against a customized GRCh38 reference genome in which the mitochondrial reference sequence was updated with the rCRS^22^. This strategy standardized mitochondrial genomic coordinates across datasets and enabled consistent heteroplasmy calling relative to the canonical mitochondrial reference used by MITOMAP (https://mitomap.org/bin/view.pl/MITOMAP/HumanMitoSeq). Mitochondrial reads were subsequently extracted from aligned BAM files with samtools^41,42^ for downstream coverage analysis and variant calling.

### scRNA-seq data processing and quality control

scRNA-seq data processing and quality control (QC) were conducted using Seurat v4^16^. Filtered feature-barcode matrices generated by Cell Ranger were used to calculate standard cell-level QC metrics, including the number of detected genes (nFeature_RNA), total UMI counts (nCount_RNA), and the percentage of transcripts derived from mitochondrial genes (percent.mt), ribosomal genes (percent.ribo), and the dominant expressed gene (Percent.Largest.Gene). To retain high-quality cells while preserving biologically diverse populations, chemistry-specific feature thresholds were applied: 200-2,500 detected genes for SC3Pv1, 200-6,000 for SC3Pv2, 400-7,000 for SC3Pv3/SC3Pv3.1, 500-10,000 for SC3Pv4, and 400-8,000 for all 5′chemistries. Additional global filtering criteria included percent.mt<75%, percent.ribo<50%, and Percent.Largest.Gene<50%. These thresholds were empirically optimized and implemented as user-configurable parameters within the WDL-based scMOCHA pipeline.

### Data normalization, dimensionality reduction and clustering

Filtered expression matrices were normalized in Seurat using SCTransform^43^, with mitochondrial (percent.mt) and ribosomal (percent.ribo) transcript percentages regressed out to mitigate technical confounding variation. Dimensionality reduction was achieved via Principal Component Analysis (PCA). The resulting principal components were utilized for graph-based clustering and parallel visualization using UMAP^44^ and t-distributed Stochastic Neighbor Embedding^45^. Cluster-specific marker genes were identified using the FindAllMarkers function via a two-sided Wilcoxon rank-sum test, restricting results to positive markers expressed in at least 10% of cells within a cluster with a minimum log_2_ fold-change threshold of 0.25 (only.pos = TRUE, min.pct = 0.1, logfc.threshold = 0.25). The top 20 marker genes per cluster, ranked by average log_2_ fold-change, were visualized using the DoHeatmap function.

### Cell type annotation

Automated cell-type annotation was performed using the reference-based mapping toolkit Azimuth^16^. Queries were matched to tissue-specific references, such as pbmcref for PBMC and bonemarrowref for BM, at a user-specified annotation granularity level (*e.g.*, celltype.l1, celltype.l2, or celltype.l3). For samples exhibiting insufficient mapping confidence or lacking a tissue-matched reference, the unsupervised, SCTransform-derived cluster identities were retained as the default cell-type labels. These finalized labels were subsequently appended as custom cluster tags (CJ) tags to corresponding alignments within the mitochondrial BAM file, facilitating downstream per-cluster pseudobulk variant calling.

### Mitochondrial BAM preparation

To generate lineage-specific datasets, cell-type metadata were integrated directly into the mitochondrial sequence alignments. Cell-type annotations were appended to corresponding cell-barcoded reads within the mitochondrial BAM file using the addtags function of the sinto toolkit (https://timoast.github.io/sinto/). This process generated a custom CJ tag for each aligned read, directly mapping it to its assigned cluster identity. The newly annotated BAM file was indexed using samtools index and subsequently partitioned by cell cluster via bamtools split using the - tag CJ parameter^46^, yielding distinct, pseudobulk BAM files for each isolated cell type. Concurrently, a bulk-labeled BAM file encompassing all retained cells was synthesized to facilitate sample-level mitochondrial haplogroup inference and global coverage profiling.

### Single-cell level and cluster level mitochondrial variant calling

Mitochondrial variants were identified at both single-cell and per-cluster pseudobulk resolutions using the mgatk framework (v0.6.2)^15^. Single-cell data were processed via mgatk tenx using filtered cell barcodes (CB), while pseudobulk data were processed via mgatk bcall using the previously appended CJ. Strand-resolved nucleotide counts (A, T, C, G) were compiled across the 16,569-bp rCRS contig. Candidate variants were defined by evaluating the set difference between observed alleles and the reference genome, and they were recorded using standard positional nomenclature (*e.g.*, 3243A>G).

For each candidate variant, cellular and cell-cluster pseudobulk AF were calculated as the ratio of alternative-allele reads (forward + reverse) to the total depth at that genomic position:

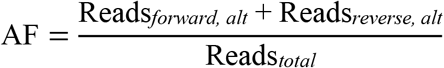

AF was assigned NA when the total read depth was <10. When the total read depth was ≥10, AF was assigned 0 unless the alternative allele was supported by at least two forward and two reverse reads at the single-cell level, or at least three forward and three reverse reads at the cell-cluster pseudobulk level.

To characterize variant distribution and reliability, we computed: (i) the variance-to-mean ratio (VMR) across the cell population, (ii) the strand-correlation coefficient between forward and reverse alternative-allele counts (excluding cells lacking coverage on both strands), (iii) the maximum observed cellular heteroplasmy, and (iv) the absolute cell counts exceeding discrete heteroplasmy thresholds of 5% and 95%. Variants were retained for downstream analysis if they were confidently detected in at least three cells (single-cell mode) or a minimum of one cluster (per-cluster mode) with an overall AF≥0.05. These empirical thresholds were selected to optimize sensitivity for low-frequency somatic heteroplasmy while stringently minimizing false positives arising from sequencing artifacts, aligning with established single-cell mitochondrial lineage-tracing.

### Haplogroup inference and variant annotation

Maternal haplogroups were assigned at the sample level using HaploGrep3^19^ based on the PhyloTree-mt 17.2 phylogenetic backbone applied to pseudobulk variant profiles. To extensively annotate variant properties including locus context, pathogenicity predictions, disease associations, and nucleotide change classifications (transitions or transversions), we queried a local SQLite snapshot of the MITOMAP database^17^. Population allele frequencies were cross-referenced against the gnomAD v3.1 mitochondrial dataset^47^. Functional impact predictions were stratified by variant genomic location: coding missense mutations were scored using APOGEE2 via MitImpact v3.1.3^23^, while non-coding elements were assessed using nAPOGEE^24^. The final comprehensive output, compiled into a standardized variant annotation table, captured: chromosome position, reference and alternative alleles, locus, associated diseases, mutation validation status, nucleotide change, amino acid substitution, evolutionary conservation (phastCons100way), MITOMAP population frequency, gnomAD population frequency, predicted computational pathogenicity tier, and the assigned haplogroup.

### mtDNA variant classification

Following the integration of single cell, pseudobulk, and global allele frequencies, each identified variant was stratified into either homoplasmic or heteroplasmic variants:

- **Homoplasmic**: For each donor, the mean AF was calculated across all cell types with a total read depth ≥10. Variants with a mean AF ≥0.95 were classified as homoplasmic.
- **Heteroplasmic**: Non homoplasmic variants.
- **Somatic**: subset of heteroplasmic variants only present in at least one but not all cell types, which satisfy both of the following criteria: (1) at least 1 cell type has sufficient number of cells (n≥10) and all cells contain only WT reads; (2) at least 1 cell type has sufficient number of cells (n≥10) that contain mutant variant (0.05≤AF≤0.95).

To safeguard against alignment artifacts, positions flagged as recurrent sequencing anomalies, including coordinates overlapping rCRS low-complexity regions, poly-N tracks, mis-alignment errors, RNA editing sites or the ENCODE hg38 genomic blacklist, were excluded prior to downstream categorical classification.

### Pre-filtering of germline and technical artifacts

Prior to somatic nomination, all tracked variants are evaluated against three independent exclusion manifests to remove inherited and technical variants:

1. Haplogroup Signatures: Lineage-defining variants are extracted from the per-sample HaploGrep3 profile based on the PhyloTree-mt 17.2 baseline and immediately purged from the somatic candidate pool.
2. Common Population Variants: Variants matching known inherited alleles are filtered using the gnomAD v3.1 mitochondrial release. Loci with a population homoplasmic allele frequency (AF_homo_) exceeding a permissive cutoff, or those found as homoplasmies in ≥5% of an in-house baseline cohort, are categorized as germline polymorphisms.
3. Reference-Track Artifacts: False positives are mitigated by filtering out variants intersecting rCRS poly-N blocks, low-complexity repeats, or the standardized ENCODE hg38 genomic blacklist, consistent with core mgatk recommendations.

### Somatic-mutation hotspot analysis

To profile cohort-wide mutational trends, the number of individuals carrying each distinct somatic variant is compiled and projected onto the 16,569-bp rCRS coordinate system using an interactive gggenes (https://wilkox.org/gggenes/) lollipop layout stratified by major structural domain (D-loop, rRNA, tRNA, and protein-coding).

### mtDNA variant heteroplasmy mapping and landscape visualization

To visualize clonal dynamics and mutation landscapes, cell-by-variant heteroplasmy matrices were sorted by cluster identity and descending summed allele frequencies. Cellular entries with a total coverage depth below 10 reads at a given variant position were systematically masked by assigning an arbitrary placeholder value of −0.1, which was rendered in grey on subsequent plots to explicitly differentiate missing data from confident reference calls. Heatmaps were generated using the ComplexHeatmap framework^48^ with a continuous viridis color scale. These matrices were augmented with row annotations denoting MITOMAP population frequency, gnomAD frequency, and haplogroup classification status, alongside column annotations tracking cell-level cluster identity, percentage of mitochondrial transcripts, and log10-transformed total UMI counts. Additionally, circular genomic plots were constructed via the circlize package^49^ to comprehensively map the genomic distribution, sample frequency (proportion of samples carrying each variant), mean pseudobulk AF, and regional density of distinct mtDNA variant classes across the entire mitochondrial genome.

### Cell culture

All cell lines were maintained at 37°C in a humidified incubator with 5% CO2. The human osteosarcoma thymidine kinase-deficient cell line 143B(TK^−^) (ATCC CRL-8303) and human embryonic kidney HEK293 cells were maintained in Dulbecco’s Modified Eagle Medium GlutaMAX (DMEM; Thermo Fisher Scientific, 10569044) containing high glucose (4.5 g/L) and sodium pyruvate (110 mg/L), supplemented with 1% non-essential amino acids (NEAA) and 10% fetal bovine serum (FBS). The human lung adenocarcinoma cell line A549 and the WAL2A lymphoblastoid^50^ cell line were cultured in RPMI 1640 medium with ATCC modifications (Thermo Fisher Scientific, A1049101), supplemented with 1% NEAA and 10% FBS. Corresponding mitochondrial DNA-depleted (ρ⁰ or Rho0) derivatives were maintained in the same medium supplemented with 50 μg/mL uridine (Sigma-Aldrich, U3750-100G). Adherent cell lines were passaged at 70-95% confluence by washing with PBS (Thermo Fisher Scientific, 14190250) and dissociating with 0.25% trypsin-EDTA (Thermo Fisher Scientific, 25200114). Cells used for comparative experiments were harvested at the same passage number.

### 10X scRNA-seq library preparation and sequencing

For the WT and Rho0 samples, a mixture of four cell lines (143B, WAL2A, HEK293, A549) was processed using the Chromium Single Cell 3′ Reagent Kit v3.1 (10X Genomics). Cell barcoding, reverse transcription, cDNA amplification, and 3′ gene expression library preparation were performed according to the manufacturer’s protocol. The resulting libraries were sequenced on either a DNBSEQ-T7 or an Illumina NovaSeq 6000 platform. The scRNA-seq data are deposited in the Gene Expression Omnibus (GEO) GSE337162.

### PacBio library preparation and sequencing

Genomic DNA was extracted from cell pellets and quantified using the Qubit dsDNA HS Assay Kit (Thermo Fisher Scientific, Q33231). Full-length mitochondrial genomes were amplified by long-range PCR using PrimeSTAR GXL DNA Premix (Takara, R051B) with custom primers and reaction conditions as previously described^51^. PCR products were purified with SMRTbell Cleanup Beads (PacBio, 102-158-300) and assessed for fragment size distribution using the Femto Pulse system (Agilent, FP-1002-0275). PacBio libraries were prepared using the SMRTbell Prep Kit 3.0 (PacBio, 102-182-700) according to their standard protocol. Following adapter ligation and nuclease treatment, libraries were size-selected using AMPure PB beads (PacBio, 100-265-900). Libraries were annealed to sequencing primers using the Sequel II Binding Kit 3.2 (PacBio, 102-333-300), bound to Sequel II Polymerase 2.2, loaded onto SMRT Cell 8M (PacBio, 102-281-700), and sequenced on a PacBio Sequel IIe platform using the Sequel II Sequencing Kit 2.0 (PacBio, 102-194-400) with a 30-hour movie collection time. The data are deposited in the Gene Expression Omnibus (GEO) GSE337162.

### PacBio long-read DNA-seq data analysis

Reads were aligned with pbmm2 to the GRCh38 reference genome, in which the mitochondrial contig (chrM) was duplicated head-to-tail (33,138 bp) to accommodate mitochondrial genome circularity; mitochondrial positions were folded to canonical 16,569 bp rCRS coordinates. Reads with mapping quality (MAPQ)≥40 and PacBio predicted read quality (rq)≥0.998 were retained. Reads in the following categories were excluded: putative NUMT-derived reads identified from supplementary alignments (≥500 bp, MAPQ≥20), reads outside the expected long-range PCR amplicon boundaries (±100 bp from primer positions), and reads<10 kb. Variant AF was calculated as the number of reads carrying the variant allele divided by the total number of reads spanning that position. Homopolymer-associated indels were excluded based on the systematic error pattern characterized in a 2.5 kb mtDNA plasmid control, with position-level assessment against the control applied within the region it covered. Variants with AF≥5% were reported.

### Correlation profiling of mitochondrial mutation burden

To evaluate potential relationships between mtDNA variant burden and confounding biological or technical variables within PBMC and BM cohorts, systematic correlation analyses were executed. The absolute counts of homoplasmic, heteroplasmic, and somatic variants determined per sample were correlated against donor age using Pearson correlation coefficients overlaid with a LOESS smoothing curve via ggpubr::ggscatter. Conversely, stratified group comparisons tracking variant burden shifts across categorical sex designations were evaluated using t tests.

### Single-variant visualization

For every prioritized variant, scMOCHA outputs a cohesive, four-panel cellular layout that groups cells horizontally to communicate data density at a glance (as shown in Fig. S6b,e,h). Ordered sequentially by variant state and descending allele frequency, the visualization stacks:

1. Cellular alternative allele frequency (AF<0.01 is masked as NA and rendered in white).
2. Local sequence depth, transformed as log_2_(depth+1).
3. The discrete four-state (variant in red, WT in blue, no sufficient reads in grey, no reads in white) evidence classification tier.
4. The corresponding single-cell cell-type label.

### Statistical assessment of cell type-specific heteroplasmy

To identify cell type-specific variations, we evaluated whether the single-cell allele frequency distribution of each detected variant differed across annotated cell types. Inter-lineage distributions were evaluated using a non-parametric Kruskal-Wallis rank-sum test restricted to high-confidence cellular variant assignments from cell types with a total read depth ≥10. Cells with an ambiguous “other” lineage label were excluded from PBMC analyses but retained for BM evaluations. Calculated P-values were structured using the broom::tidy framework and adjusted for multiple testing across all variants within an individual sample using the Benjamini–Hochberg false discovery rate (FDR) procedure^52^. Variants with an adjusted P<0.05 were considered significant cell-type-specific candidates. Major cell type specific variants were selected based on the Kruskal-Wallis rank-sum test and visualized using dot plots.

### Comparative analysis of tissue-specific variant distributions

To evaluate whether mtDNA variant distributions differed between PBMC and BM microenvironments, two-sided Fisher’s exact tests were performed on non-haplogroup variants within each classified category. These statistical comparisons were stratified separately across homoplasmic, heteroplasmic, and somatic mutation cohorts. This stratified approach was utilized to uncover potential tissue-enriched or niche-specific mutational landscapes and clonal expansion profiles across hematopoietic compartments.

### Analysis of strand-specific variant distributions

To investigate strand-specific mutational asymmetry, variants were classified based on their localization on the heavy (H) or light (L) strand of the mitochondrial genome. The absolute proportions and distinct nucleotide substitution profiles (*e.g.*, specific transition and transversion frequencies) on each strand were summarized and systematically compared among homoplasmic, heteroplasmic, and somatic variant classes, with downstream plotting modules rendering the relative frequencies and strand breakdowns as structured, multi-panel bar plots.

### Characterization of mitochondrial base substitution spectra

To evaluate mutational signatures across different variant classes and tissue types, single-nucleotide variants (SNVs) were classified into the six standard pyrimidine-faced base substitution categories: C:G → T:A, C:G → A:T, C:G → G:C, T:A → C:G, T:A → G:C, and T:A → A:T, collapsing complementary mutations. The relative frequencies and compositional proportions of these six substitution types were calculated independently within each analytic stratum, crossing the three variant categories (homoplasmic, heteroplasmic, and somatic) with the two sample microenvironments (PBMC and BM). The resulting mutation profile breakdowns were visualized using stratified pie charts to display the characteristic mutational signatures of each distinct variant class.

### Single-cell multi-sample integration and variant-stratified differential expression

To assess downstream functional impacts of specific mutations, single-cell transcriptomes from all samples carrying a target variant were merged and co-embedded. Multi-sample batch effects were mitigated by performing anchor-based canonical correlation analysis (CCA) integration via the IntegrateLayers function (method = CCAIntegration) across the top 30 principal components within the Seurat v5 framework^34^. Following integration, discrete assay slots were consolidated across batches using the JoinLayers function. Within each annotated cell lineage, single cells were stratified into distinct phenotypic cohorts: a “heteroplasmy-high” alternative-positive group (AF≥0.5) and a “sufficient-reads” reference-only group (AF<0.5), bounded by cell-level mutation frequency and read depth thresholds. Finally, differential gene expression (DGE) between these groups was calculated using a two-sided Wilcoxon rank-sum test via FindMarkers, evaluating the log-normalized counts of the core RNA assay.

### Functional enrichment analysis

To characterize the downstream biological consequences of somatic mutations, significantly perturbed transcripts were analyzed using comprehensive pathway enrichment frameworks. DEGs meeting strict significance and magnitude criteria (adjusted P<0.05, |avg_log2FC|>0.25, pct.1 or pct.2≥0.05) were stratified by their direction of regulation (up- or down-regulated). To ensure proper identifier mapping of mitochondrially encoded features across annotation networks, the canonical MT-prefix was programmatically stripped from all gene symbols prior to submission. Gene Ontology (GO) Biological Processes was executed using the enrichGO function within the clusterProfiler package^53^ against the org.Hs.eg.db human genome annotation database. Statistical significance across ontologies was visualized using customized bar plots and correlation plots charting −log10-transformed adjusted P-values against individual pathway, restricted to the top 10-20 most enriched terms per category.

### Disease cohort comparisons

To evaluate differences between condition cohorts (*e.g.*, COVID-19 versus healthy controls), comparative statistical analyses were conducted across three distinct resolution tiers:

- Sample-level variant burden: Global differences in total variant burden per individual were compared using a two-sided Wilcoxon rank-sum test.
- Allele frequency distributions: Per-variant allele frequency profiles across cohorts were evaluated using a Kruskal-Wallis test, followed by Dunn’s post-hoc test for pairwise comparisons where applicable.
- Cell-type-stratified heteroplasmy: Inter-cohort variations in lineage-specific heteroplasmy were modeled using linear mixed-effects models (lme4 framework), treating condition as a fixed effect and incorporating sample-level random intercepts to control for intra-subject covariance when biological replicates were available.

### Inflammatory module score

Per-cell inflammatory gene module scores were computed on the log-normalized RNA assay of the merged Seurat object using *Seurat::AddModuleScore()* in Seurat v5^34^, which calculates, for each cell, the mean expression of the input gene set minus the mean expression of a randomly sampled control gene set binned by average expression. The inflammatory gene set comprised 30 curated genes spanning four functional categories: (i) cytokines/chemokines (*IL6*, *IL1B*, *TNF*, *CXCL8*, *CCL2*, *CCL3*, *CCL4*, *CXCL1*, *CXCL2*, *CXCL3*, *CXCL10*), (ii) NF-κB pathway components and targets (*NFKBIA*, *NFKB1*, *REL*, *RELA*), (iii) type I interferon-stimulated genes (*ISG15*, *ISG20*, *MX1*, *MX2*, *OAS1*, *IFIT1*, *IFIT2*, *IFIT3*, *IRF1*, *STAT1*), and (iv) immediate-early stress/acute-phase response genes (*FOS*, *JUN*, *EGR1*, *ATF3*, *SOCS3*, *TNFAIP3*). Before scoring, the gene set was intersected with the genes detected in the merged Seurat object; gene sets retaining fewer than three genes were excluded. *AddModuleScore* was invoked with default binning (*nbin* = 24) and a control gene-set size of *ctrl* = min(100, 5 × *n_features*), with a fixed random seed for reproducibility.

### Structural prediction and analysis of 8362T>G (MT-TK)

For the 8362T>G variant located in the mitochondrial tRNA region, secondary RNA structures of both WT and variant sequences were predicted using RNA secondary structure prediction tool RNAstructure^54^. The structure with the minimum free energy was selected as the final model. Three-dimensional RNA structures were subsequently generated using RNAComposer^55^ based on the predicted secondary structures. Structural visualization and comparison between WT and MUT models were performed using PyMOL (V3.1.0) to evaluate mutation-associated conformational alterations and structural rearrangements.

### Structural modeling and molecular analysis of 11736T>C (*MT-ND4*, p.L326P) and 11232T>C (*MT-ND4*, p.L158P)

Structural analyses were conducted to investigate the potential functional effects of the 11736T>C (ND4, L326P) and 11232T>C (ND4, L158P) variants. Mutant protein sequences were generated by introducing target amino acid substitutions directly into the wild-type mitochondrial protein sequences. Three-dimensional structures of the wild-type protein were obtained from the Protein Data Bank (PDB; Complex I structure: 9I4I), while mutant structures were predicted using AlphaFold3^56^. Structural alignment, superposition, and visualization were subsequently performed using PyMOL to evaluate mutation-associated conformational changes. Local hydrogen-bonding interactions surrounding the mutation sites were analyzed and compared between the wild-type and mutant structures. To quantify local conformational disruptions, angular measurements were performed between helical segments flanking the L326/P326 position for 11736T>C, and a similar geometric analysis was applied to the 11232T>C variant.

### Evolutionary conservation analysis of *MT-TK* 8362T

Nucleotide sequences corresponding to mitochondrial tRNA-Lys (*MT-TK*) from representative vertebrate and invertebrate species were retrieved from the NCBI nucleotide database using the accession numbers indicated in the figure (Fig. 3b). Multiple sequence alignment was performed using MAFFT with default parameters and visualized in Jalview^57^. Conserved residues surrounding the human *MT-TK* 8362T position were manually inspected across species. Sequence logo visualization was generated using WebLogo to summarize nucleotide conservation within the aligned region.

### Evolutionary conservation analysis of ND4 L326

Protein sequences corresponding to mitochondrial NADH dehydrogenase subunit 4 (ND4) from representative vertebrate and invertebrate species were retrieved from the UniProt database using the accession numbers indicated in the figure (Shown in Fig. 4a). Multiple sequence alignment was performed using Clustal Omega^58^ with default parameters and visualized in Jalview^59^. Conserved residues surrounding the ND4 L326 position were manually inspected across species. Sequence logo visualization was generated using WebLogo^60^ to summarize residue conservation within the aligned region

### Statistical analysis

All statistical analyses were executed within the R computational environment. Unless otherwise specified, inter-group comparisons were performed using two-sided non-parametric frameworks: the Wilcoxon rank-sum test for two-group evaluations and the Kruskal-Wallis test for multi-group partitions. Associations between continuous variables were evaluated using parallel Pearson and Spearman rank correlation coefficients. To control for type I error inflation from multiple hypothesis testing, raw P-values were adjusted within each analytic stratum using the Benjamini– Hochberg false discovery rate (FDR) procedure. No statistical methods were applied to predetermine sample sizes; all experimental samples and individual cells satisfying our primary quality control metrics were included in downstream evaluations without further exclusion.

## Code and data availability

The scMOCHA pipeline source code, WDL definitions, conda environment configurations, and core analytical scripts are publicly available on GitHub at https://github.com/Xinglab/scMOCHA. All downstream analysis scripts and custom plotting code utilized to generate the figures presented in this manuscript are hosted at https://github.com/Xinglab/scMOCHA-data. Public single-cell RNA sequencing (scRNA-seq) datasets re-analyzed in this study could be retrieved from the NCBI GEO database and are accessible via the secondary accession numbers compiled in Supplementary Table 1.

## Acknowledgments

This project is primarily supported by the NIH Human BioMolecular Atlas Program (HuBMAP) through award U01HL166058 to L.P., D.C.W., and Y.X. L.P. is also supported by the Office of the Assistant Secretary of Defense for Health Affairs through the Peer Reviewed Medical Research Program under Awards W81XWH22-1-0058, NIH R01DK111495, U54HL165442, U24AT013504 and American Heart Association Established Investigator Award. D.C.W. is also supported by NIH awards R01AG078814 and R01 CA259635 and DOD award W81XWH2110128 PR202887. Opinions, interpretations, conclusions, and recommendations are those of the author(s) and are not necessarily endorsed by The Assistant Secretary of Defense for Health Affairs or endorsed by the Department of Defense.

## Author contributions

C.L. developed the framework and code of scMOCHA under the supervision of Y.X., and T.P. helped refine it. C.L. and T.P. used scMOCHA to analyze publicly available scRNA-seq data. P.L. performed scRNA-seq library preparation of the mixed cell lines, Z.C., P.P. and G.W. performed cell culture and PacBio bulk long-read DNA-seq library preparation of individual cell line, and T.P., P.H., Z.C. and S.Z. analyzed relevant data. M.L. advised on pathogenic significance of mtDNA variants. C.L., T.P. and L.P. wrote, and all other authors reviewed and edited the manuscript. D.C.W., Y.X. and L.P. designed and supervised the overall project and generated funding.

---

**Fig. S1.**
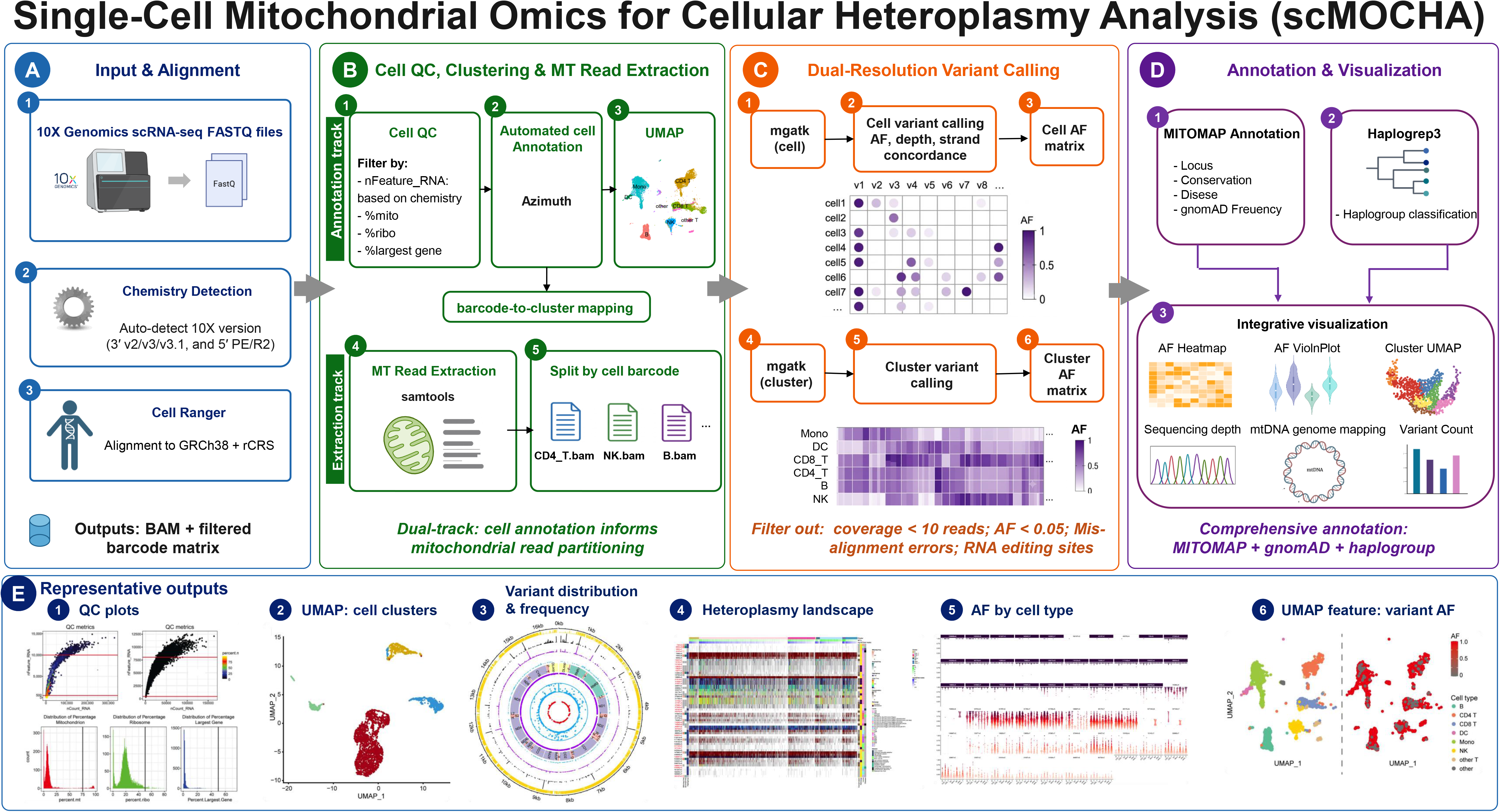
scMOCHA framework and workflow.

**Fig. S2.**
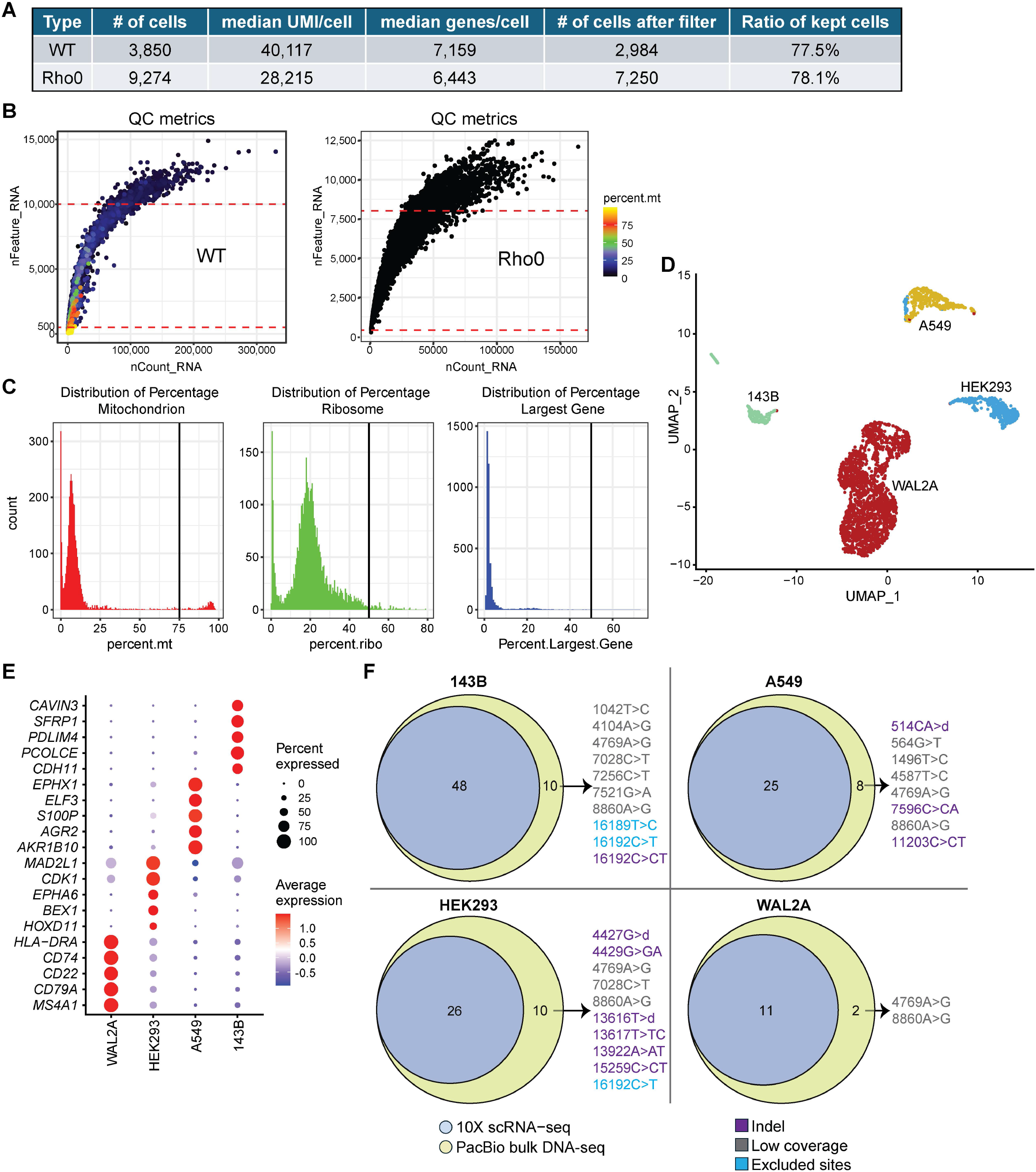
Validating mtDNA variant calling by scMOCHA. (A) Summary of scRNA-seq statistics. (B) Representative scatter plots. Filtering criteria are shown with red dashed lines. (C) Representative distribution plots showing additional filtering criteria: percent. mt<75, percent. ribo<50, and percent.Largest.Gene<50. (D-E) UMAP (D) showing 4 cell types (A549, 143B, HEK293, and WAL2A) based on gene expression profiles (E). (F) Venn diagrams comparing mtDNA variants detected by scMOCHA using scRNA-seq data (light blue) and by PacBio bulk long-read DNA-seq (light yellow) in individual cell line. Number of shared and platform-specific variants detected are shown. Variants detected only by PacBio are further classified and color-coded by reasons.

**Fig. S3.**
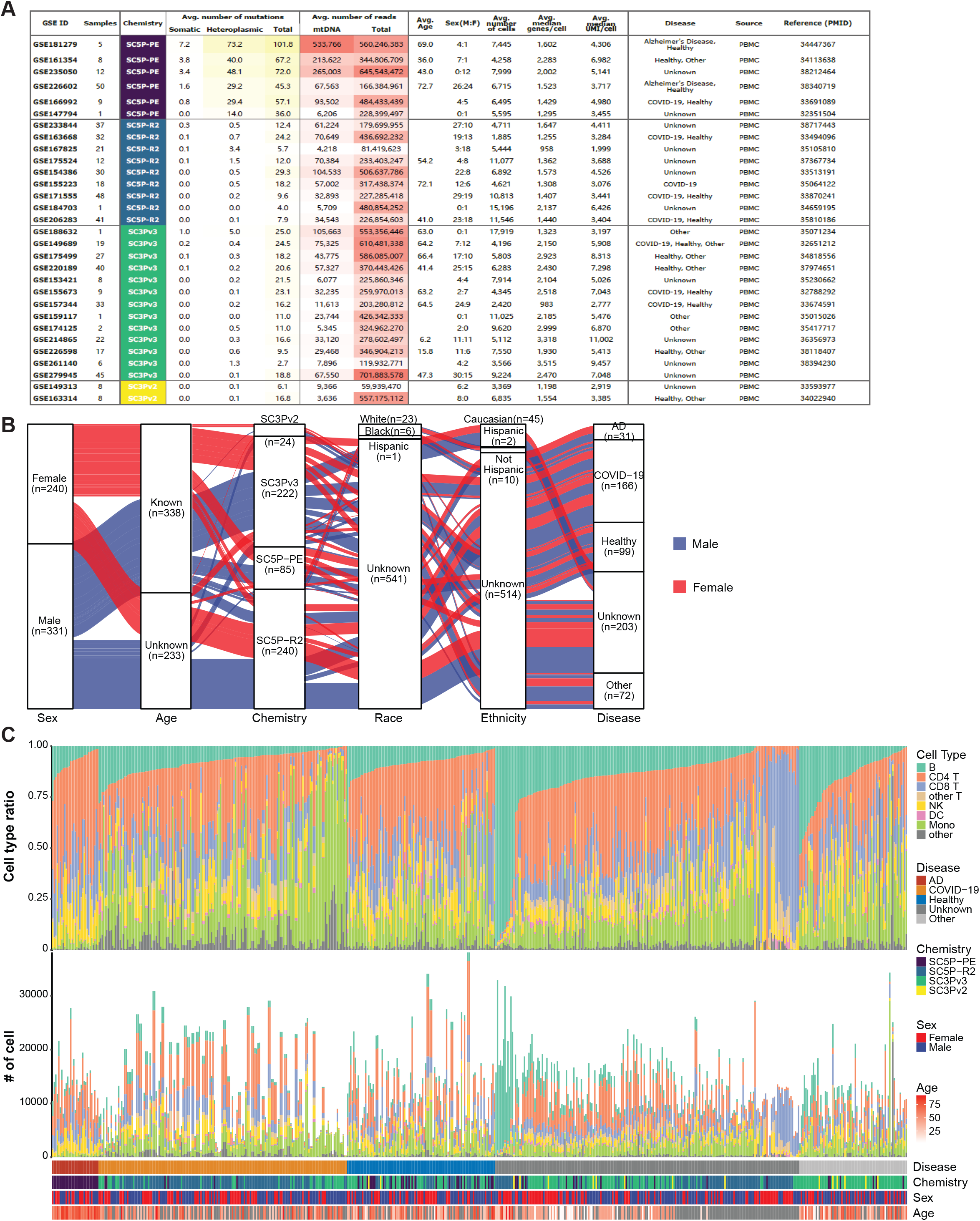
Overview of the PBMC scRNA-seq datasets. (A) Summary table of all datasets analyzed. (B) Alluvial plot showing the distribution across various sample metadata. AD - Alzheimer’s Disease. “Other” diseases include Hemorrhagic Fever with Renal Syndrome, Inflammatory Bowel Disease, Influenza, STING-Associated Vasculopathy with Onset in Infancy, and Severe Fever with Thrombocytopenia Syndrome. (C) Top: stacked bar plot showing the relative cell-type composition of each sample. Colors indicate annotated cell types. Bottom: Stacked bar plot showing the absolute number of cells per cell type for each sample. Sample-level metadata are shown as colored annotation tracks beneath the plot. For age, gray indicates samples with missing age information.

**Fig. S4.**
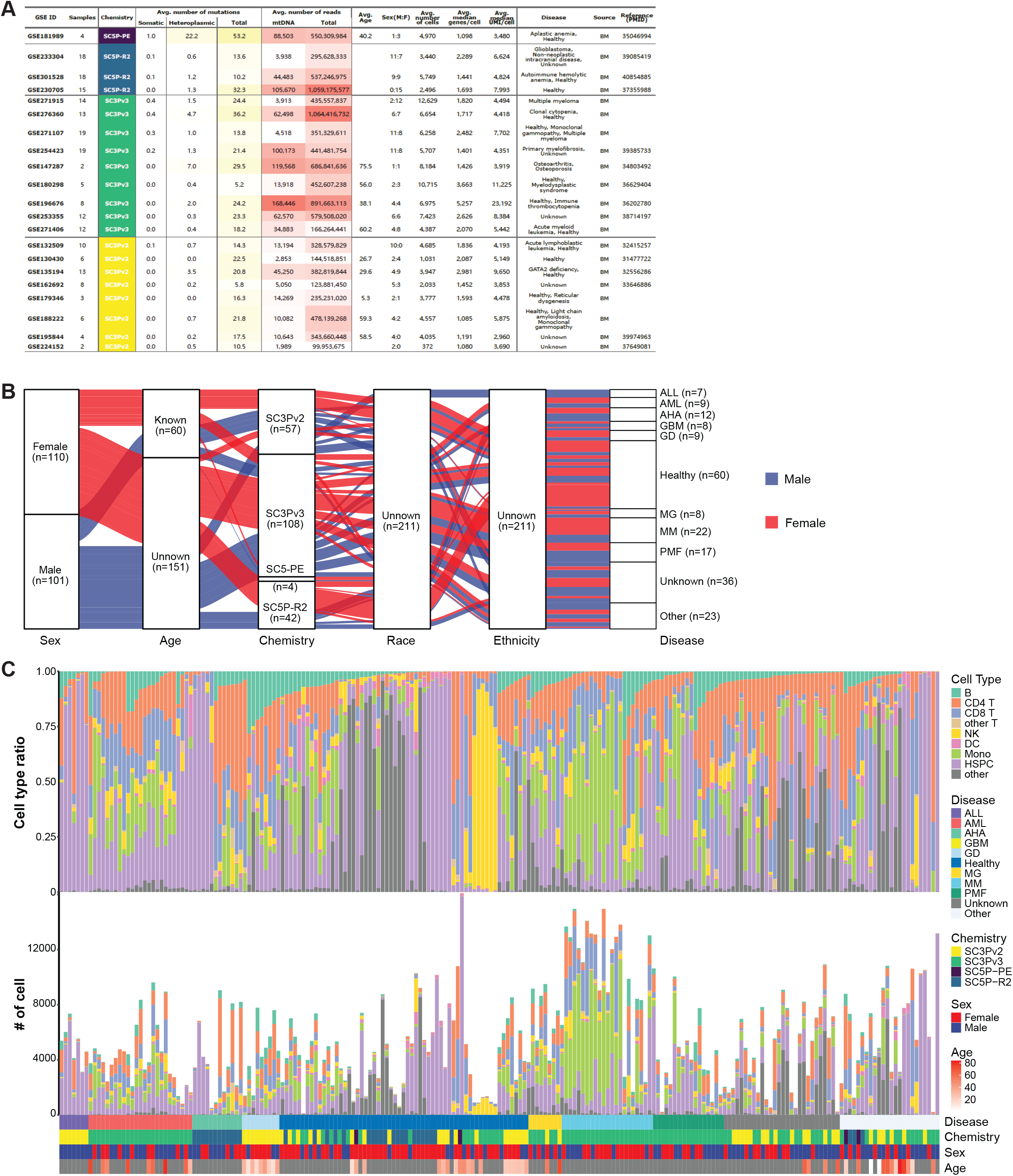
Overview of the bone marrow (BM) scRNA-seq datasets included in this study. (A) Summary table of all datasets analyzed. (B) Alluvial plot showing the distribution across various sample metadata. (C) Top: stacked bar plot showing the relative cell-type composition of each sample, with colors indicating annotated cell types. Bottom: stacked bar plot showing the absolute number of cells per cell type for each sample. Sample-level metadata are displayed as colored annotation tracks beneath the plot. For age, gray indicates samples with missing age information. Abbreviations: AHA - autoimmune hemolytic anemia; ALL - acute lymphoblastic leukemia; AML - acute myeloid leukemia; GBM - glioblastoma; GD - GATA2 deficiency; MG - monoclonal gammopathy; MM - multiple myeloma; PMF - primary myelofibrosis. “Other” diseases in (B) include aplastic anemia, clonal cytopenia of undetermined significance, immune thrombocytopenia, light chain amyloidosis, myelodysplastic syndrome, reticular dysgenesis, osteoarthritis, and osteoporosis.

**Fig. S5.**
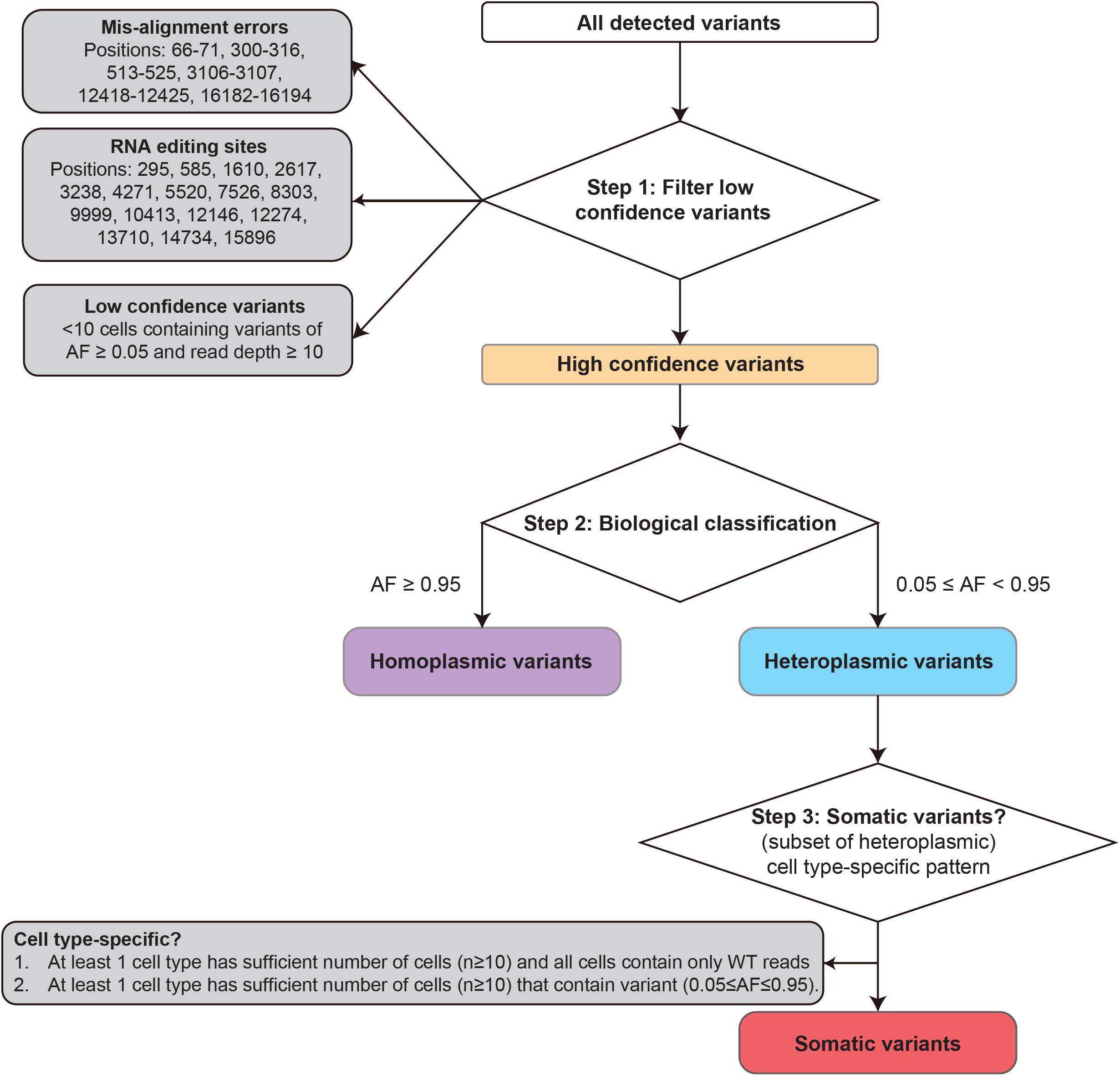
Variant classification flow chart and parameters in scMOCHA. AF: allele frequency.

**Fig. S6.**
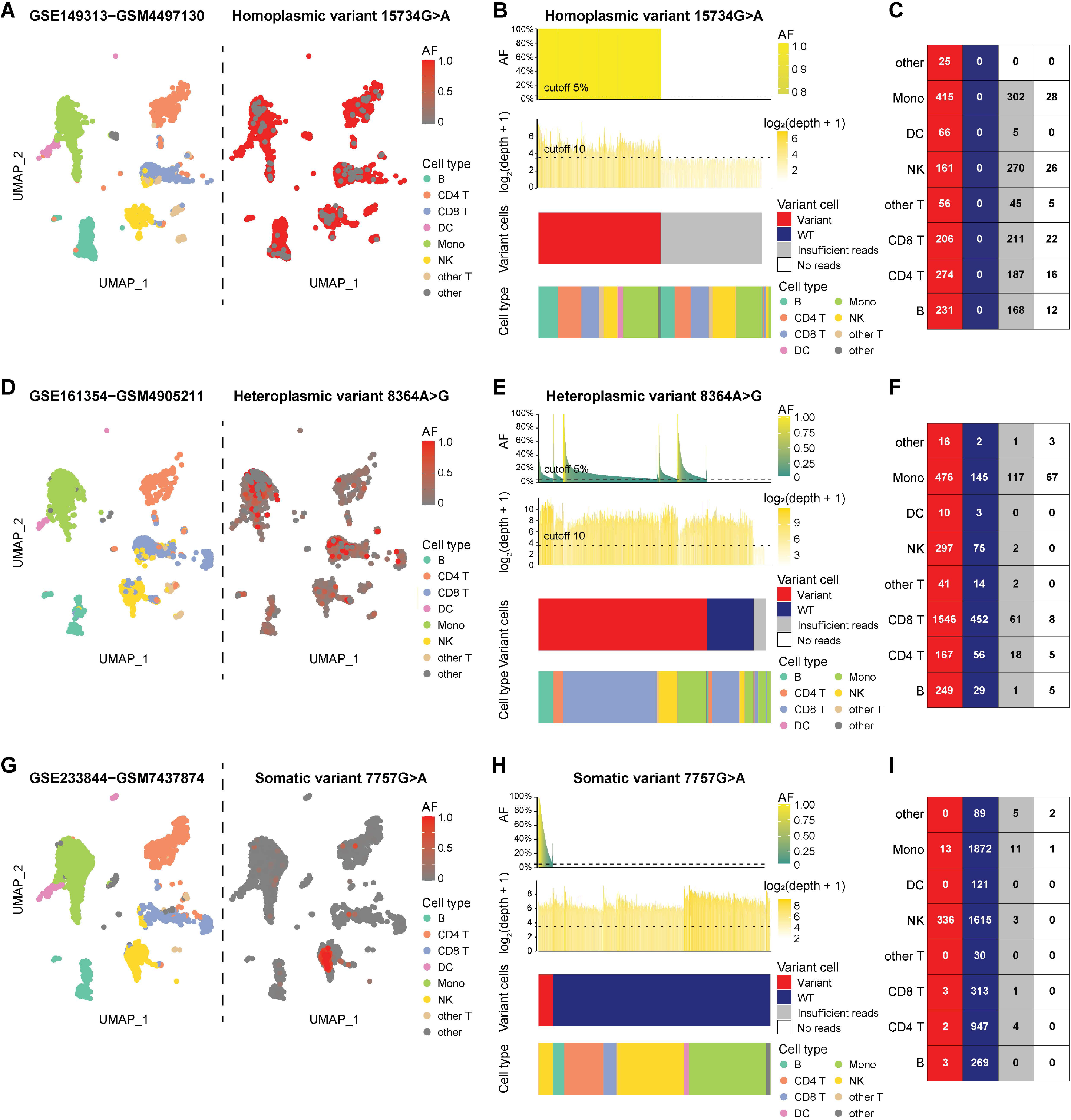
Examples of homoplasmic (A-C), heteroplasmic (D-F), and somatic (H-l) mtDNA variants. (A, D, G) UMAPs of the PBMC dataset showing the cell types (left) and variant cell distribution (right). (B, E, H) From top to bottom: variant AF, sequencing read depth, variant containing cells, and cell types. (C, F, I) Number of variant containing cells at single-cell level.

**Fig. S7.**
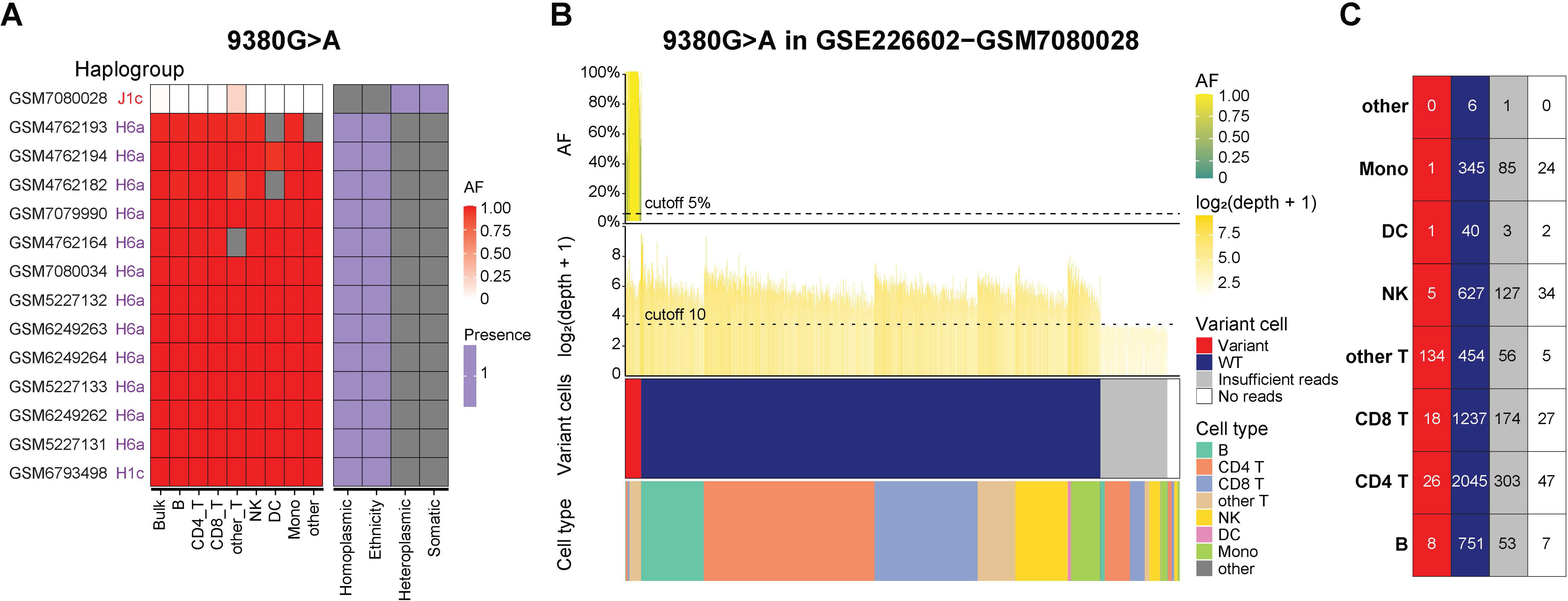
Example of a variant that is homoplasmic for some donors (all those in H1c or H6a haplogroups) but heteroplasmic and somatic for anther donor (J1c, GSM7080028). (A) mtDNA haplogroup and cell type allele frequency (AF) at pseudobulk levels are shown. (B) Single-cell AF, sequencing read depth, cell number for different variant classification, and cell type annotation for each individual cell. (C) Cell counts classified as variant-containing (red), wild-type (blue), insufficient sequencing coverage (gray), or no sequencing coverage (white).

**Fig. S8.**
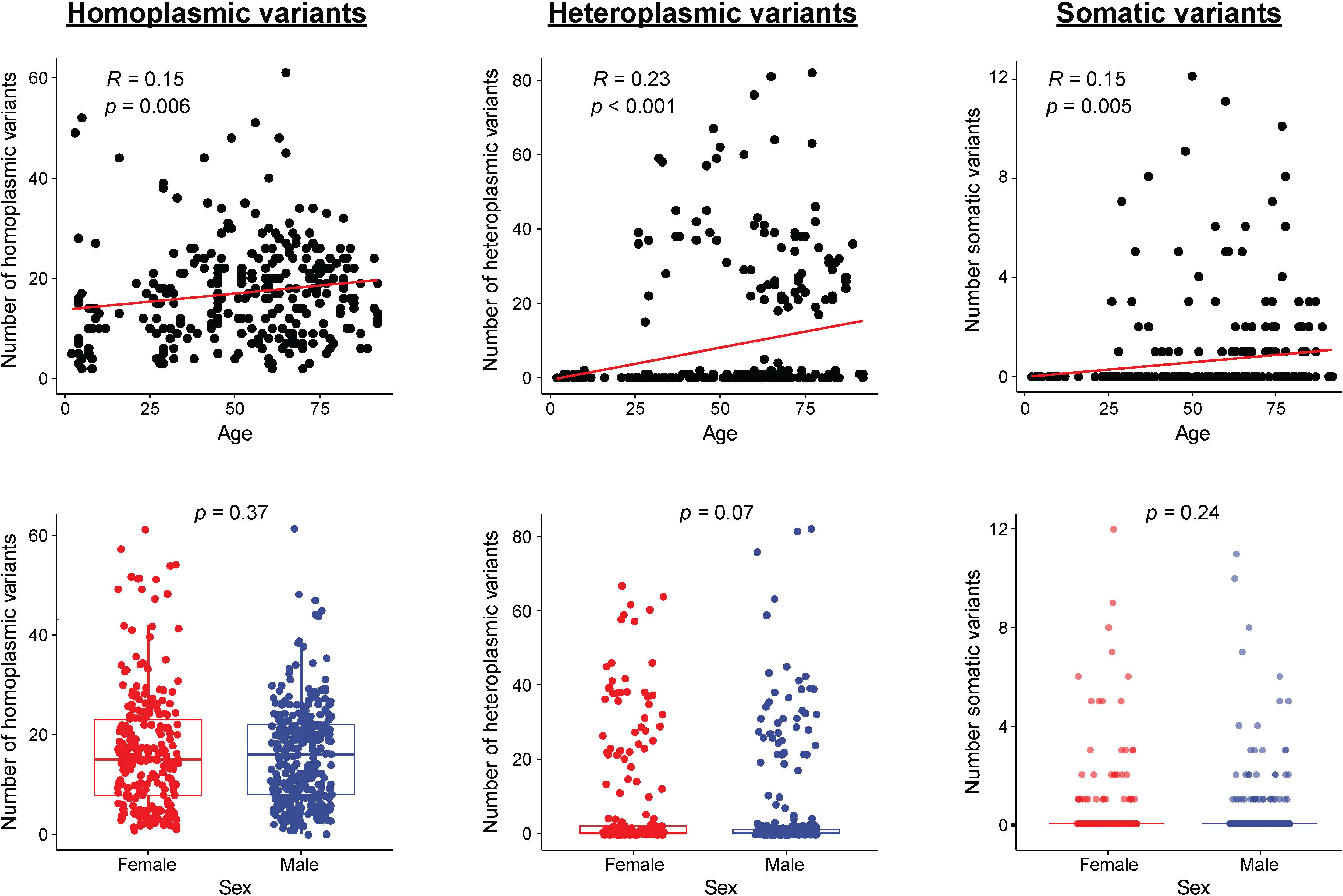
Correlation of number of homoplasmic, heteroplasmic and somatic mtDNA variants with. age and sex in PBMC samples.

**Fig. S9.**
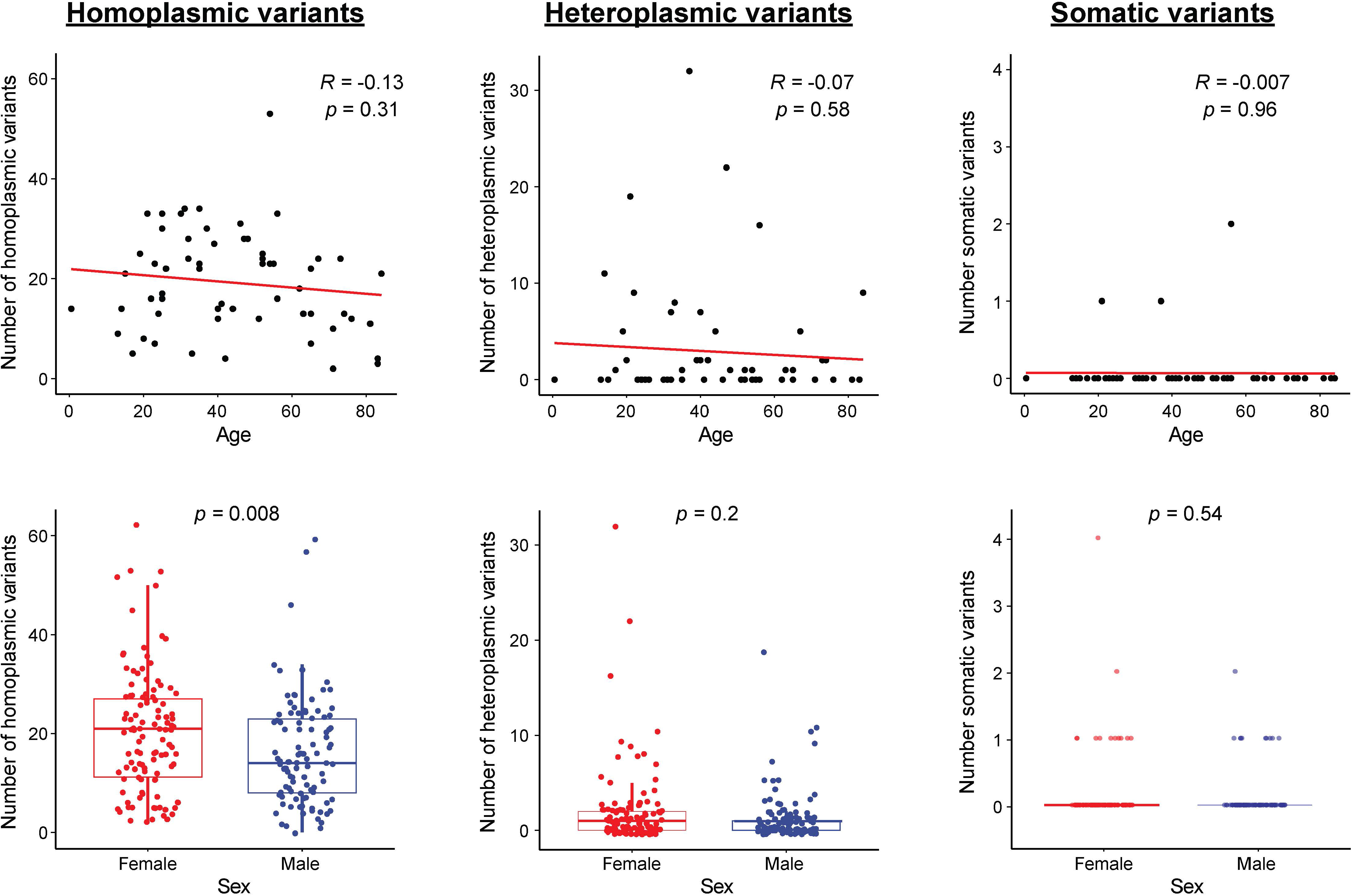
Correlation of number of homoplasmic, heteroplasmic and somatic mtDNA variants with age and sex in BM samples.

**Fig. S10.**
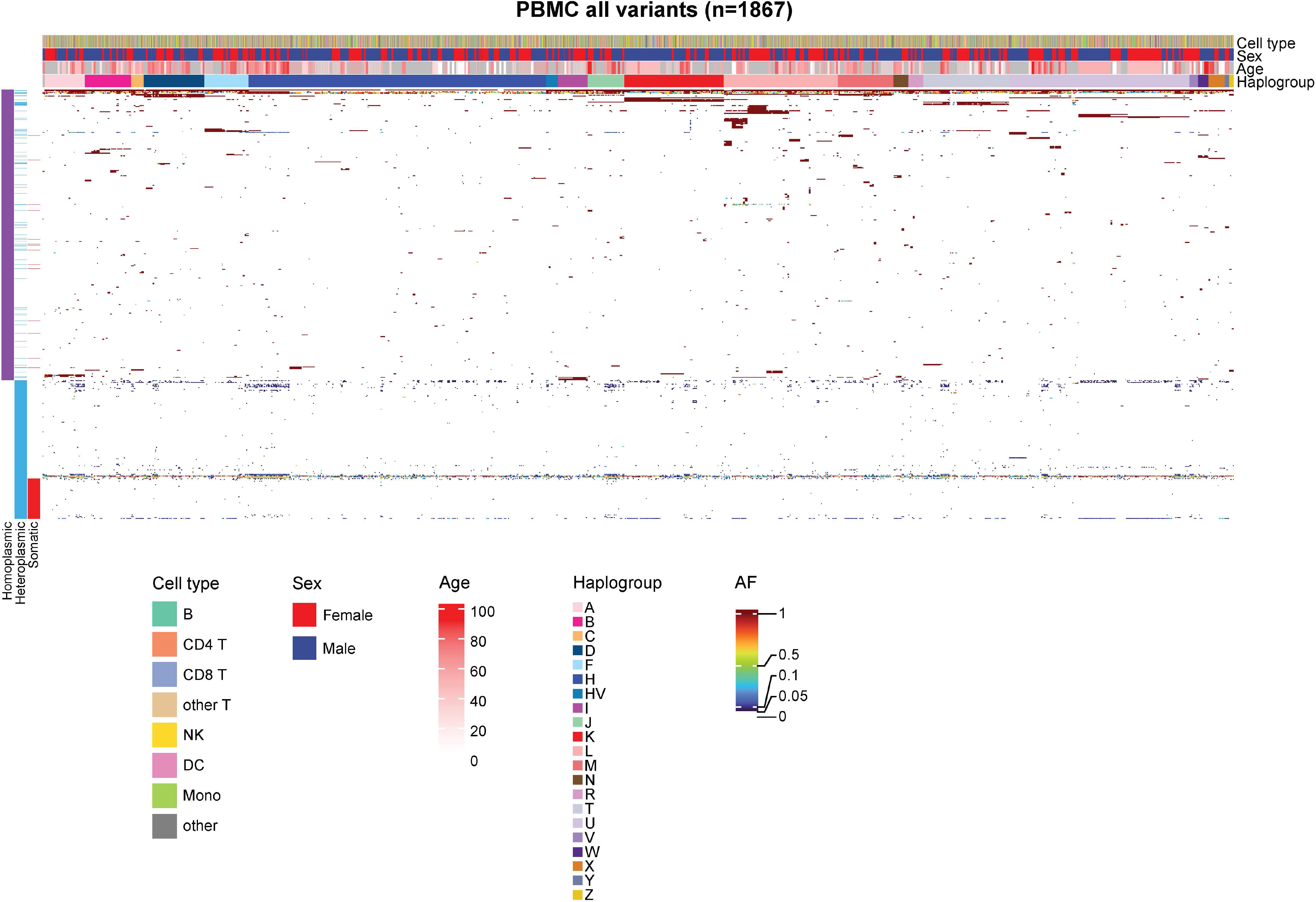
Landscape of PBMC mtDNA variants. AF - allele frequency.

**Fig. S11.**
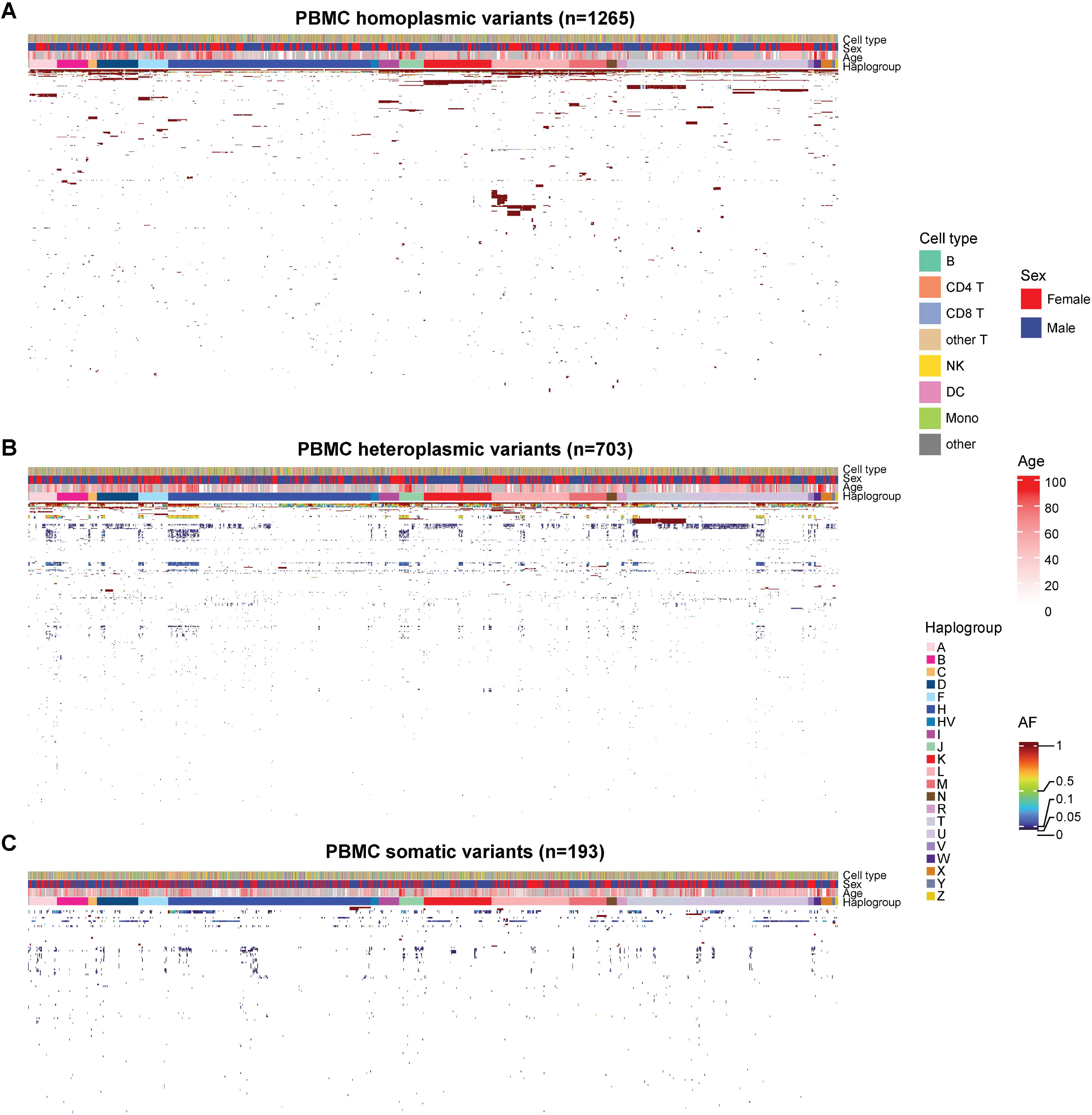
Landscape of PBMC mtDNA homoplasmic (A), heteroplsamic (B) and somatic (C) variants. AF - allele frequency.

**Fig. S12.**
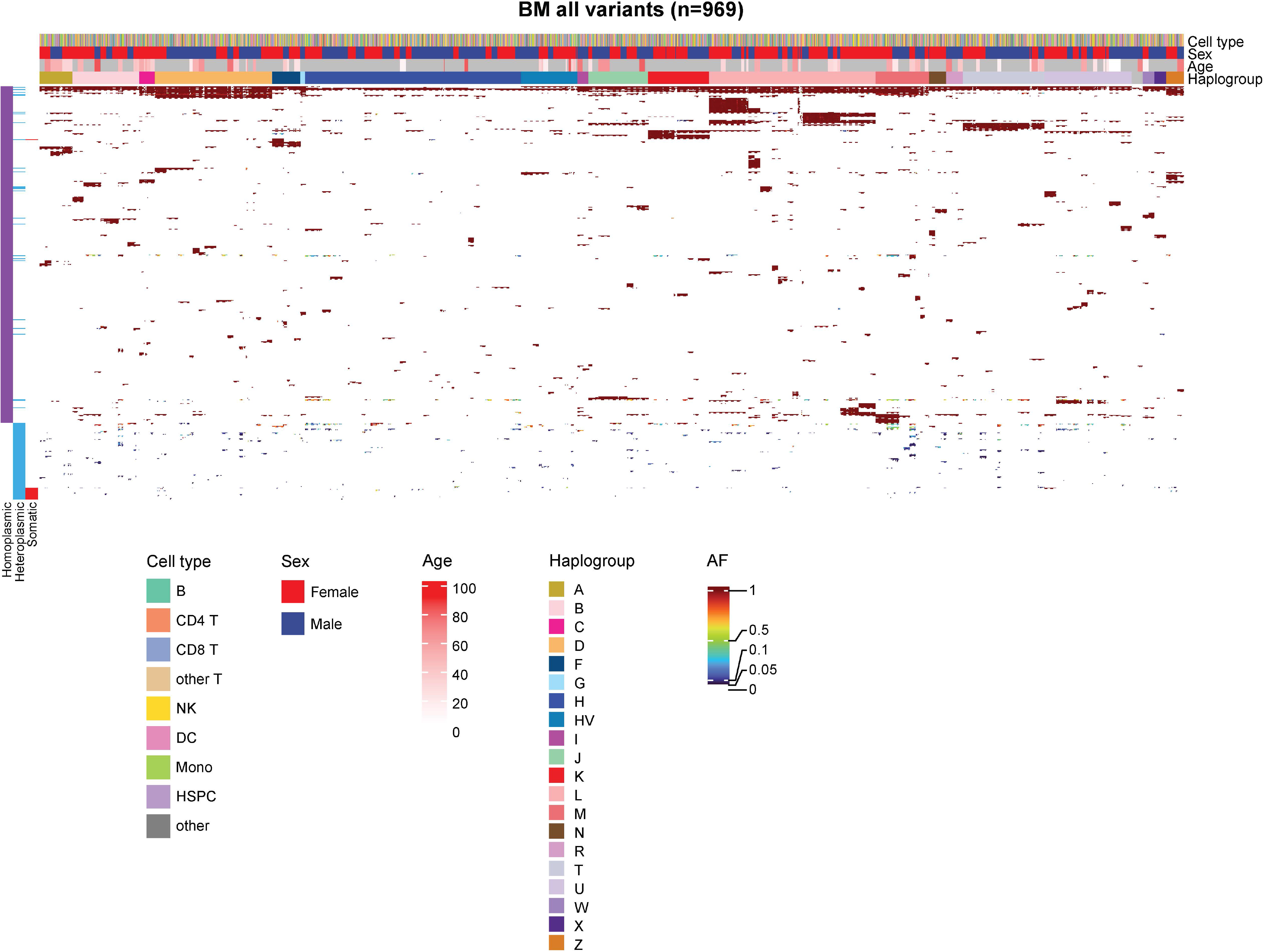
Landscape of BM mtDNA variants. AF - allele frequency.

**Fig. S13.**
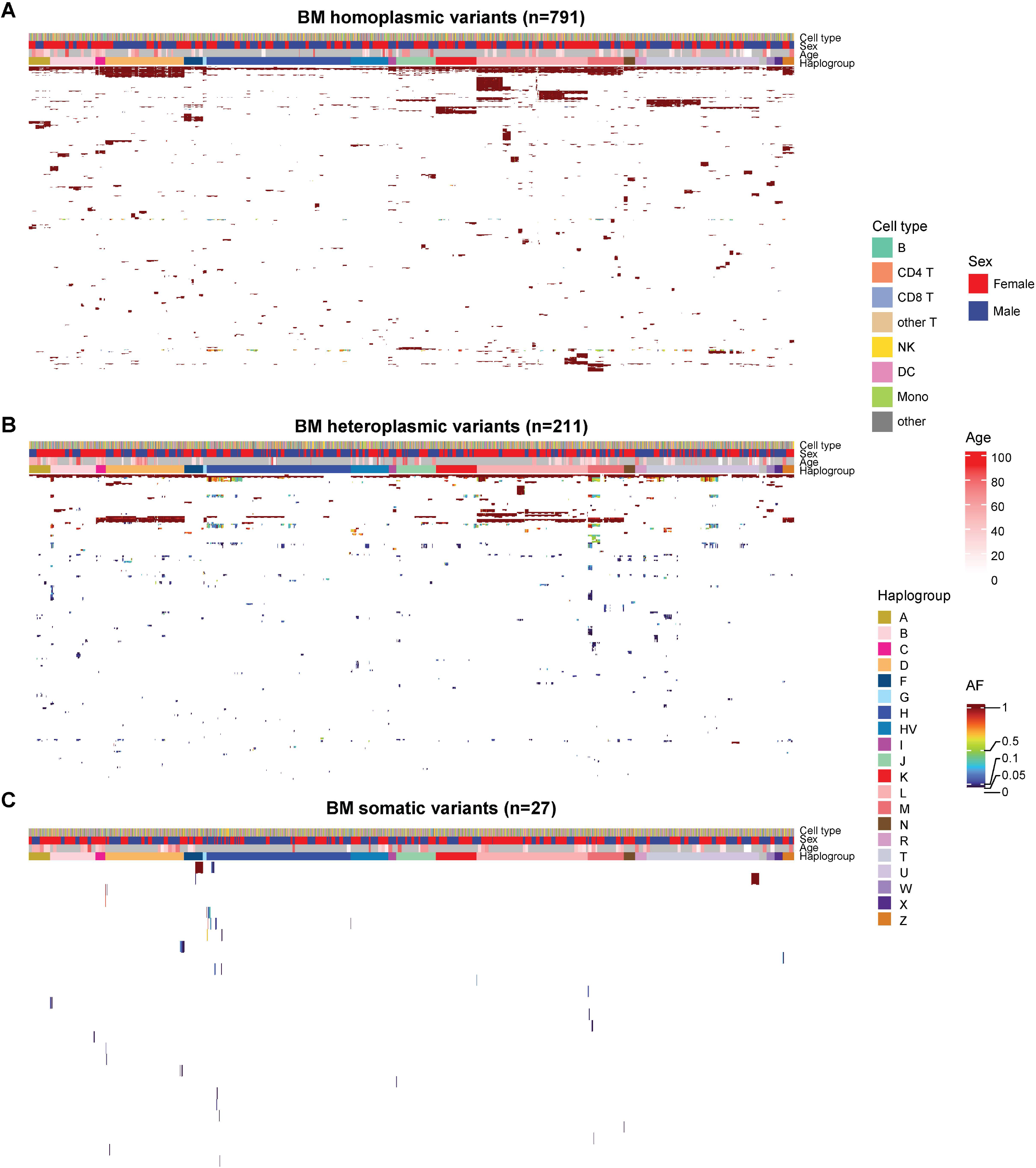
Landscape of BM mtDNA homoplasmic (A), heteroplsamic (B) and somatic (C) variants. AF - allele frequency.

**Fig. S14.**
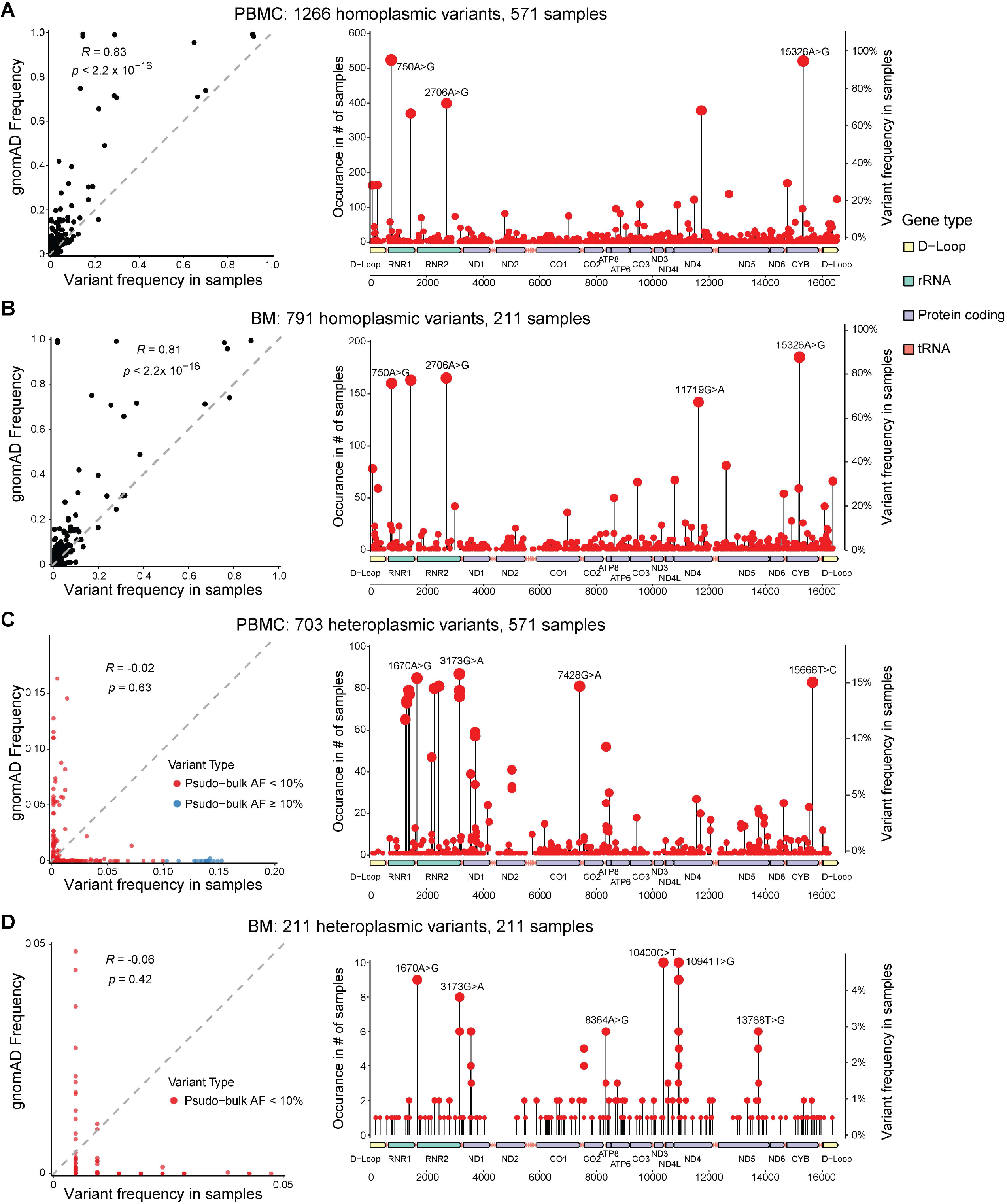
Correlation between variant frenquency of all samples of this study and their corresponding frequencies in gnomAD (left), and distribution of mtDNA variants across the mitochondrial genome with their occurrence in the study cohort (right). (A) Homoplasmic variants in PBMC. (B) Homoplasmic variants in BM. (C) Heteroplasmic variants in PBMC. (D) Heteroplasmic variants in BM. Each dot represents a unique variant.

**Fig. S15.**
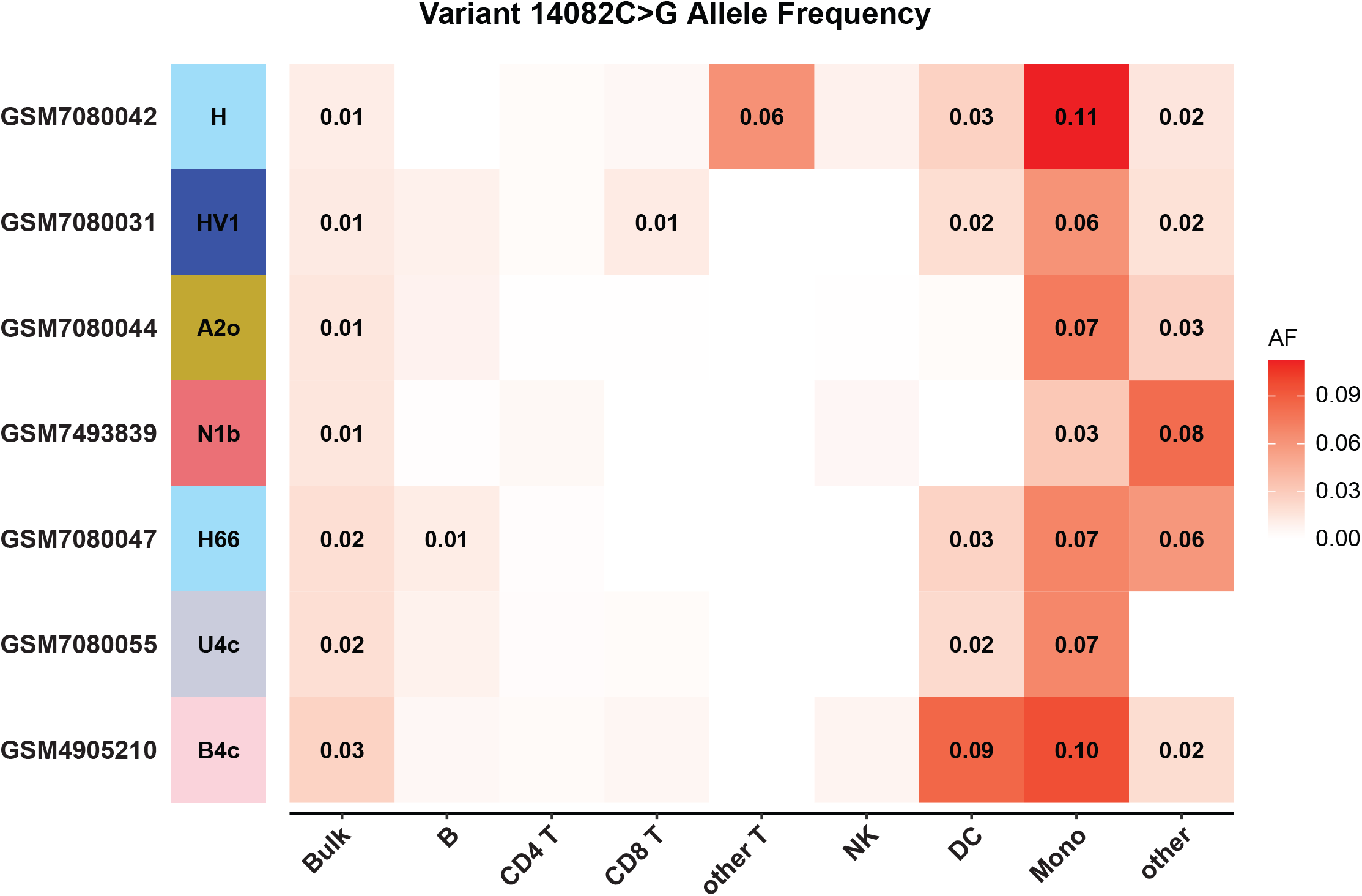
Pseudobulk and cell type-specific heteroplasmy of PBMC hotspot variant 14082C>G.

**Fig. S16.**
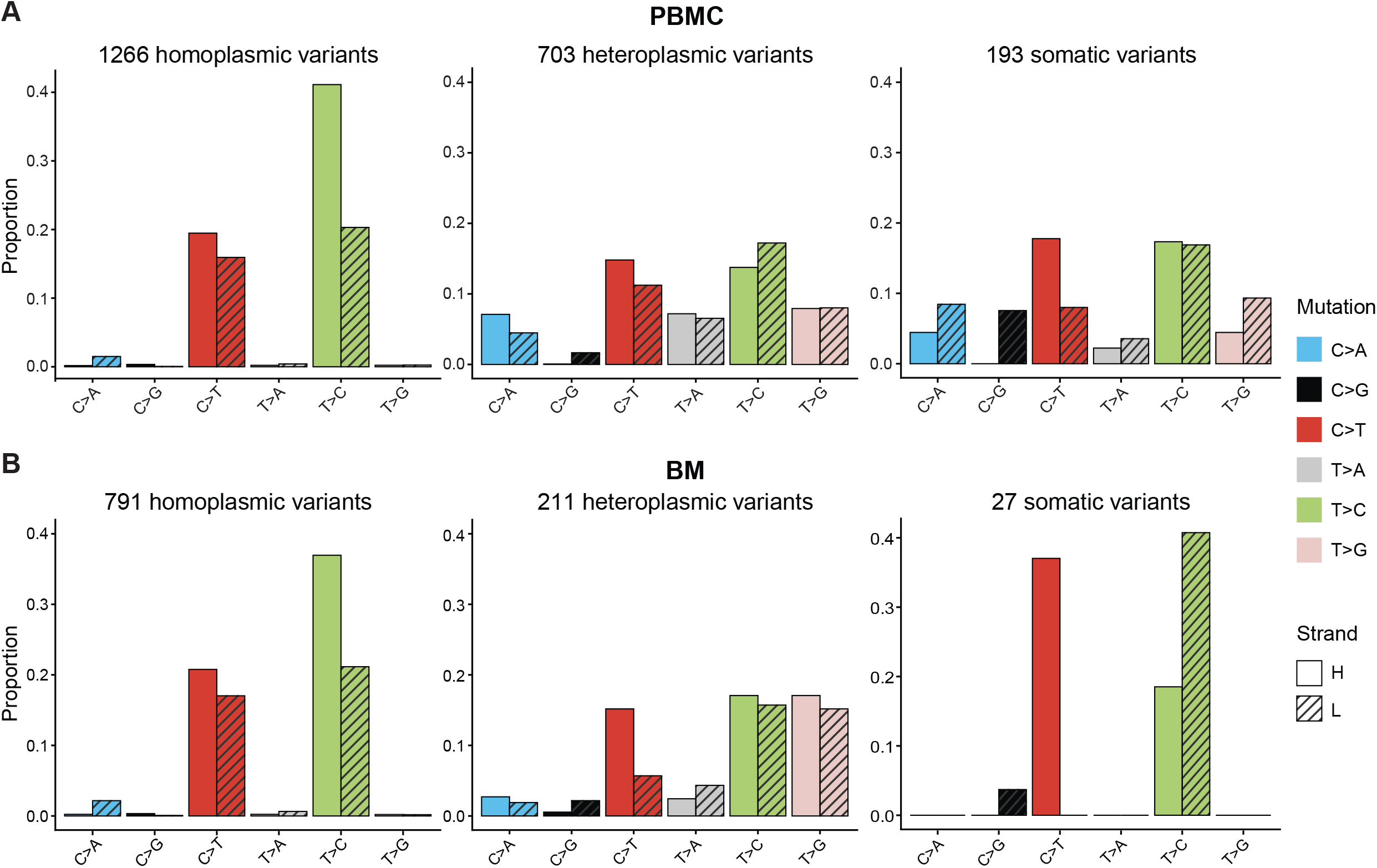
Distribution of mutation types among homoplasmic (left), heteroplasmic (middle), and somatic variants (right) in PBMC (A) and BM (B). Bar plots show the proportion of each substitution type (C>A, C>G, C>T, T>A, T>C, T>G), with strand orientation indicated as heavy (H) and light (L). Colors represent mutation classes as indicated in the legend.

**Fig. S17.**
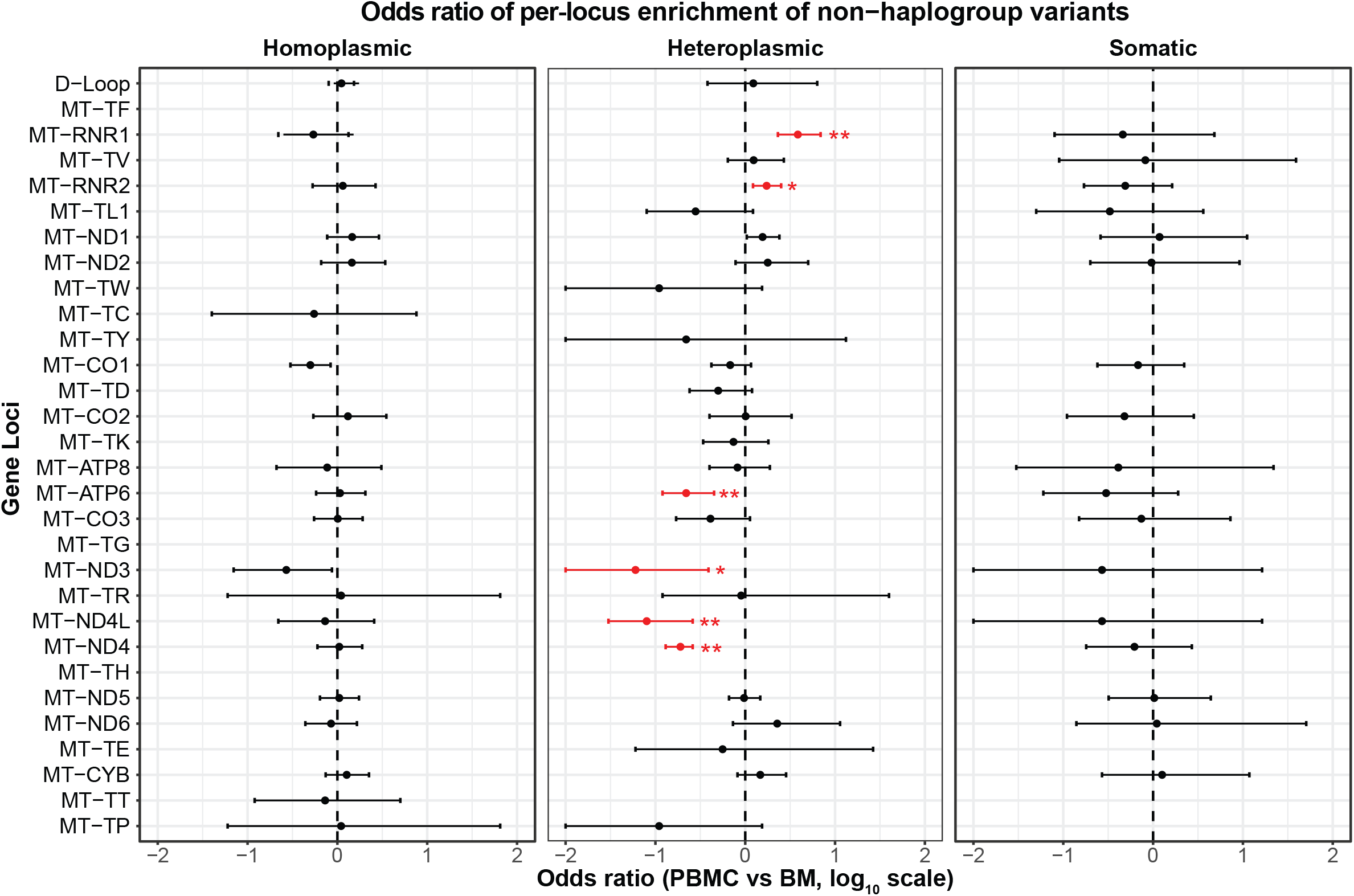
Per-locus enrichment of non-haplogroup mtDNA variants between PBMC and BM. Forest plots showing odds ratios (log,_10_scale) for per-locus enrichment of non-haplogroup variants in PBMC relative to BM across mitochondrial genomic regions. Results are shown separately for homoplasmic (left), heteroplasmic (middle), and somatic variants (right). Each point represents the estimated odds ratio for a given locus, with horizontal lines indicating 95% confidence intervals. The dashed vertical line indicates no enrichment (log,_10_ OR = 0). Gene loci are labeled along the y-axis, including the D-loop, rRNA genes, tRNA genes, and protein-coding genes. Statistically significant enrichments are highlighted in red. * p<0.01; ** p<0.0001.

**Fig. S18.**
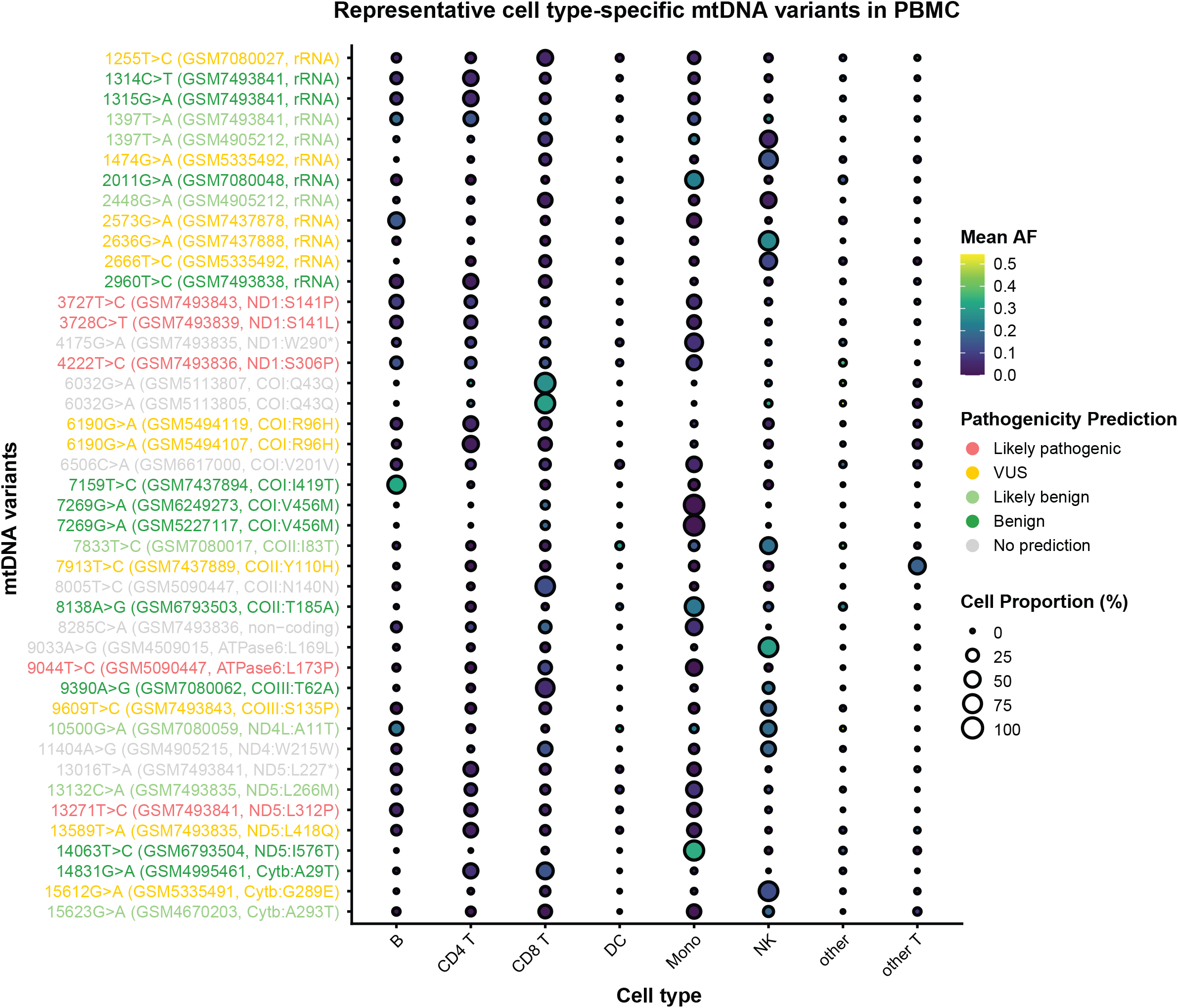
Representative cell type-specific mtDNA variants identified in PBMC samples. Dot plot showing the distribution of representative cell type-enriched mtDNA variants across major PBMC populations, including B cells, CD4 T cells, CD8 T cells, dendritic cells (DC), monocytes (Mono), natural killer (NK) cells, and other cell type. Dot color represents the mean heteroplasmy allele frequency (AF) of each variant within the corresponding cell type, while dot size indicates the proportion of cells carrying the variant. Variant labels are colored according to pathogenicity prediction categories, including likely pathogenic, variants of uncertain significance (VUS), likely benign, benign, and variants without available prediction.

**Fig. S19.**
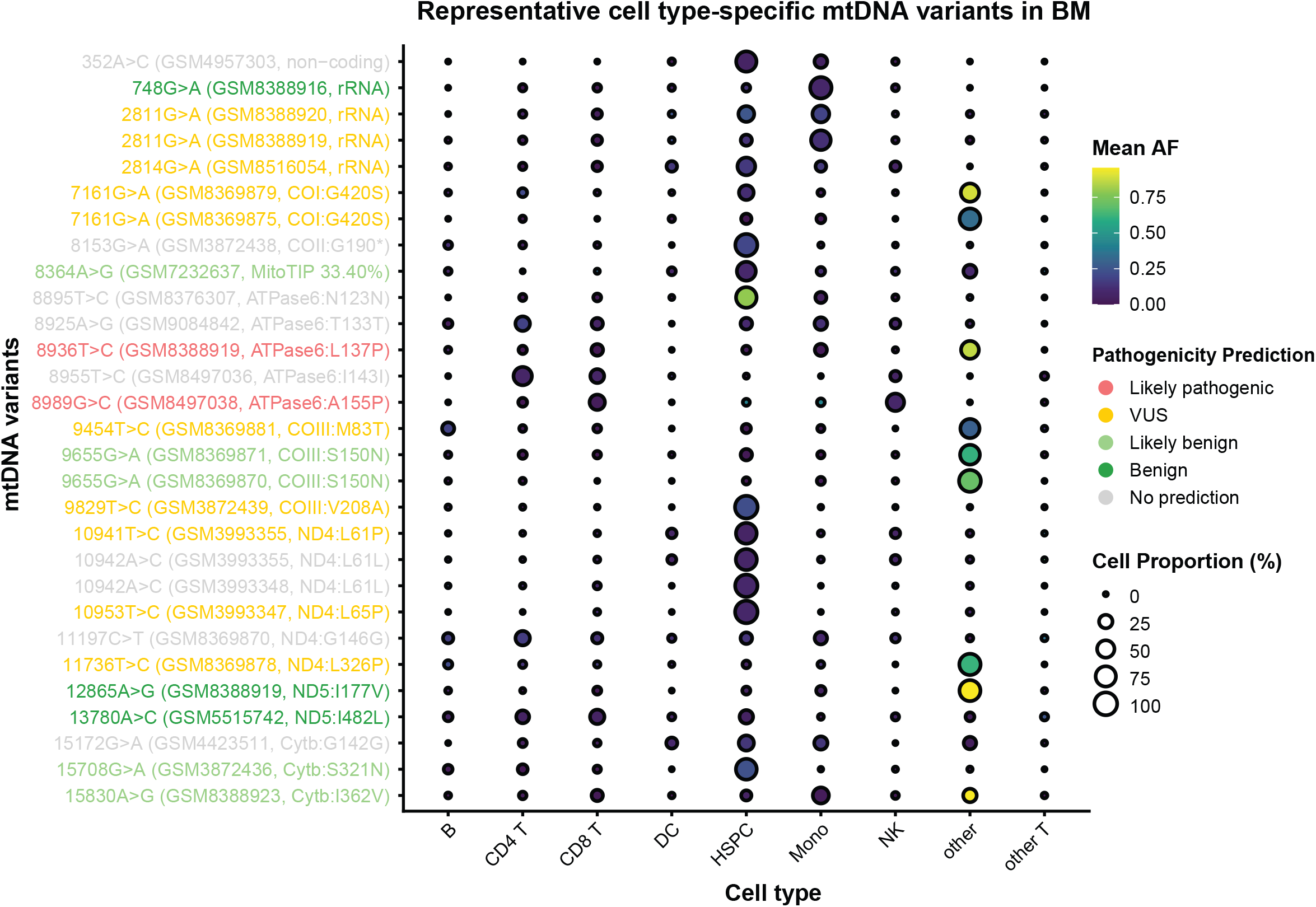
Representative cell type-specific mtDNA variants identified in BM samples. Dot plots show the distribution of representative cell type-enriched mtDNA variants across major bone marrow (BM) cell populations. In addition to major cell types of PBMC samples, hematopoietic stem and progenitor cells (HSPCs) appear in the BM.

**Fig. S20.**
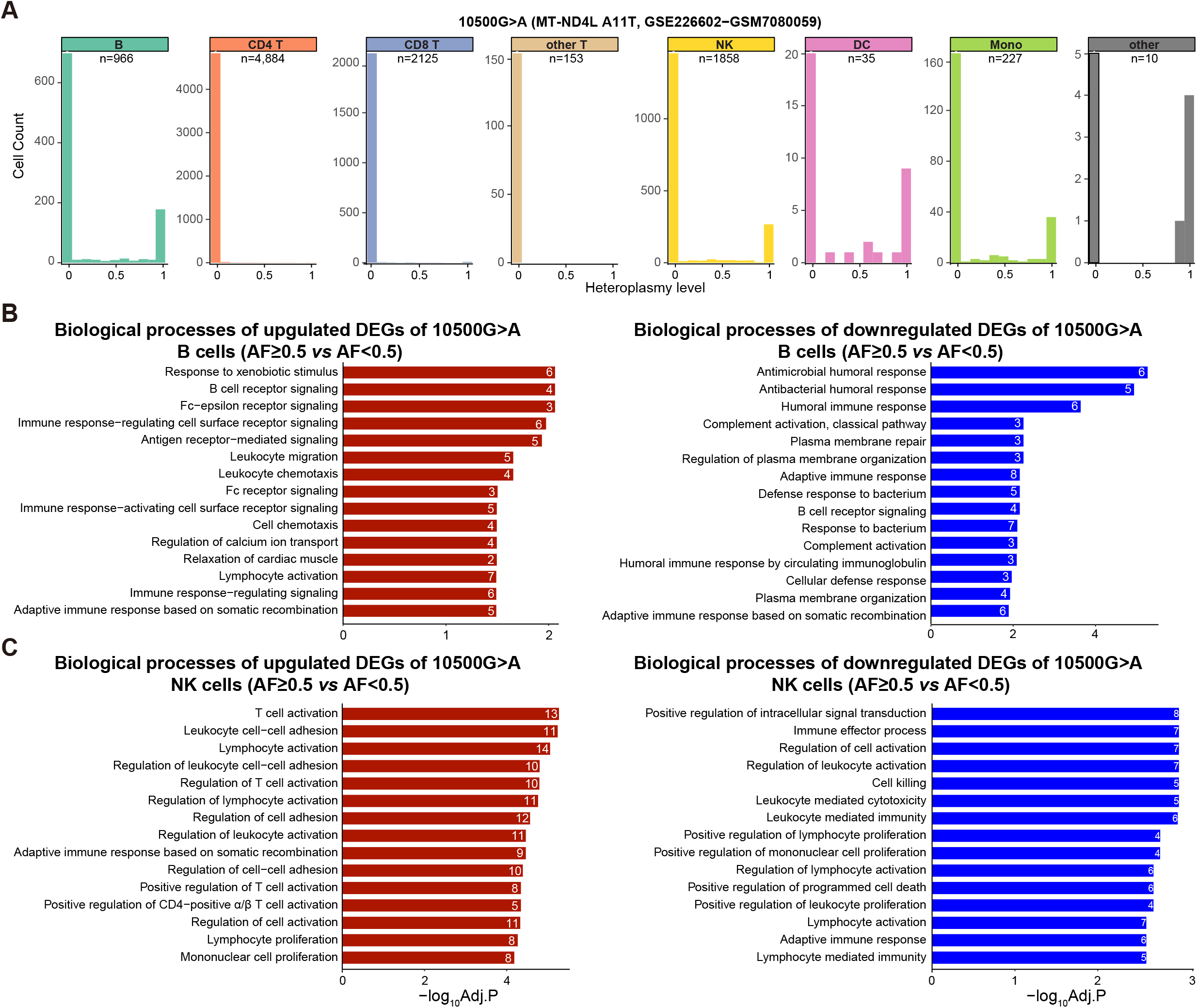
Cell type-specific distribution and transcriptional programs associated with the 10500G>A variant in PBMC. (A) Distribution of 10500G>A heteroplasmy levels across major PBMC cell populations, including B cells, CD4 T cells, CD8 T cells, NK cells, dendritic cells (DC), monocytes (Mono), and other hematopoietic cell types. The number of cells analyzed for each population is indicated above each panel. (B-C) Gene Ontology biological process (GO-BP) enrichment analyses of upregulated and downregulated differentially expressed genes (DEGs) between B cells (B) and NK cells (C) carrying high (AF≥0.5) or low (AF<0.5) heteroplasmy levels of the 10500G>A variant. Bar length represents enrichment significance as −log_10_ adjusted p-value, and numbers inside each bar indicate the number of genes associated with the corresponding biological process.

**Fig. S21.**
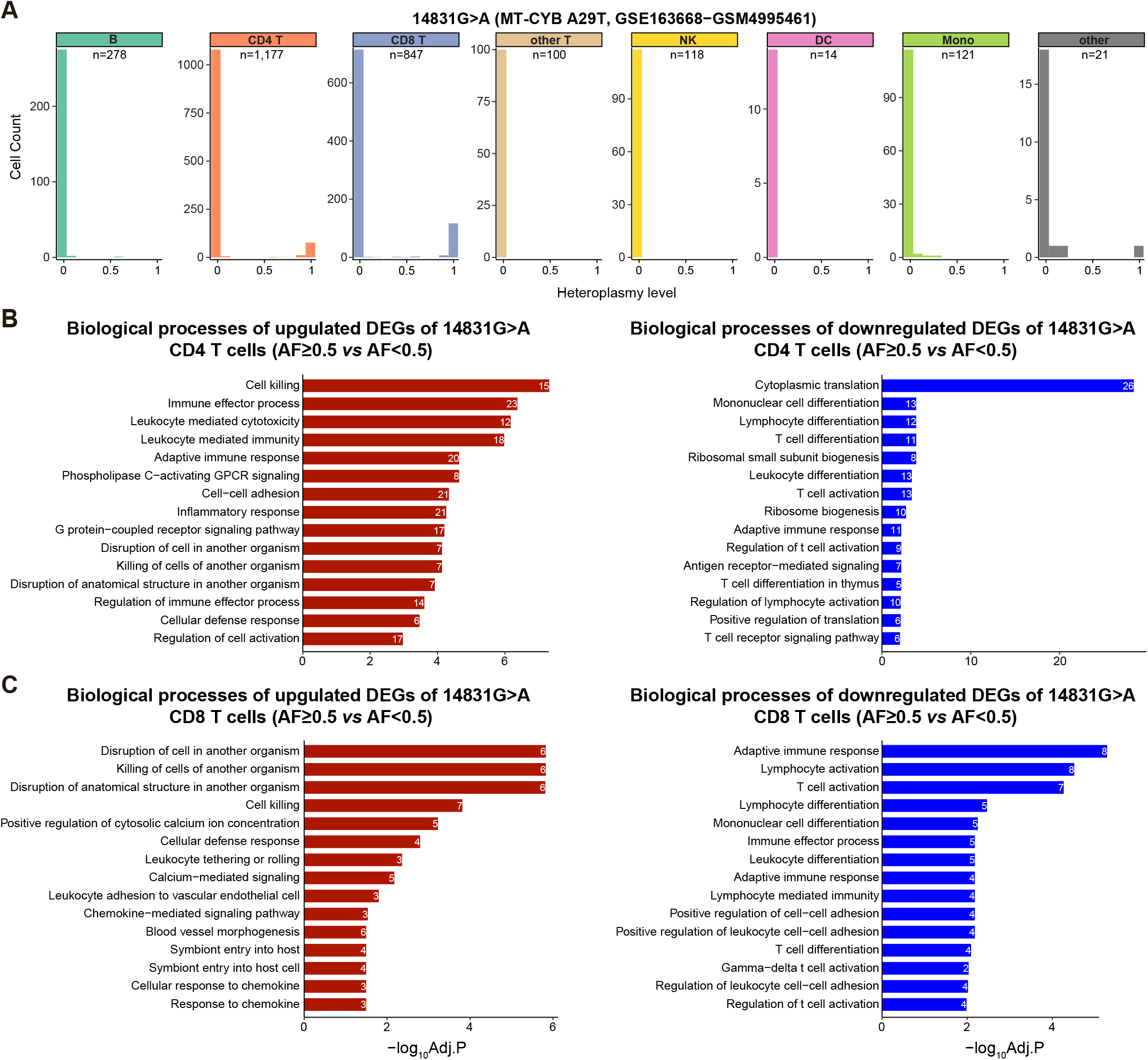
Cell type-specific distribution and transcriptional consequences of the 14831G>A variant in PBMC. (A) Distribution of heteroplasmy allele frequencies (AFs) of the 14831G>A variant across major PBMC cell populations. (B-C) GO-BP enrichment analyses of upregulated and downregulated differentially expressed genes (DEGs) between CD4 T cells (B) and CD8 T cells (C) carrying high (AF≥0.5) or low (AF<0.5) heteroplasmy levels of the 14831G>A variant. Other figure settings and annotations are the same to those described in Fig. S20.

**Fig. S22.**
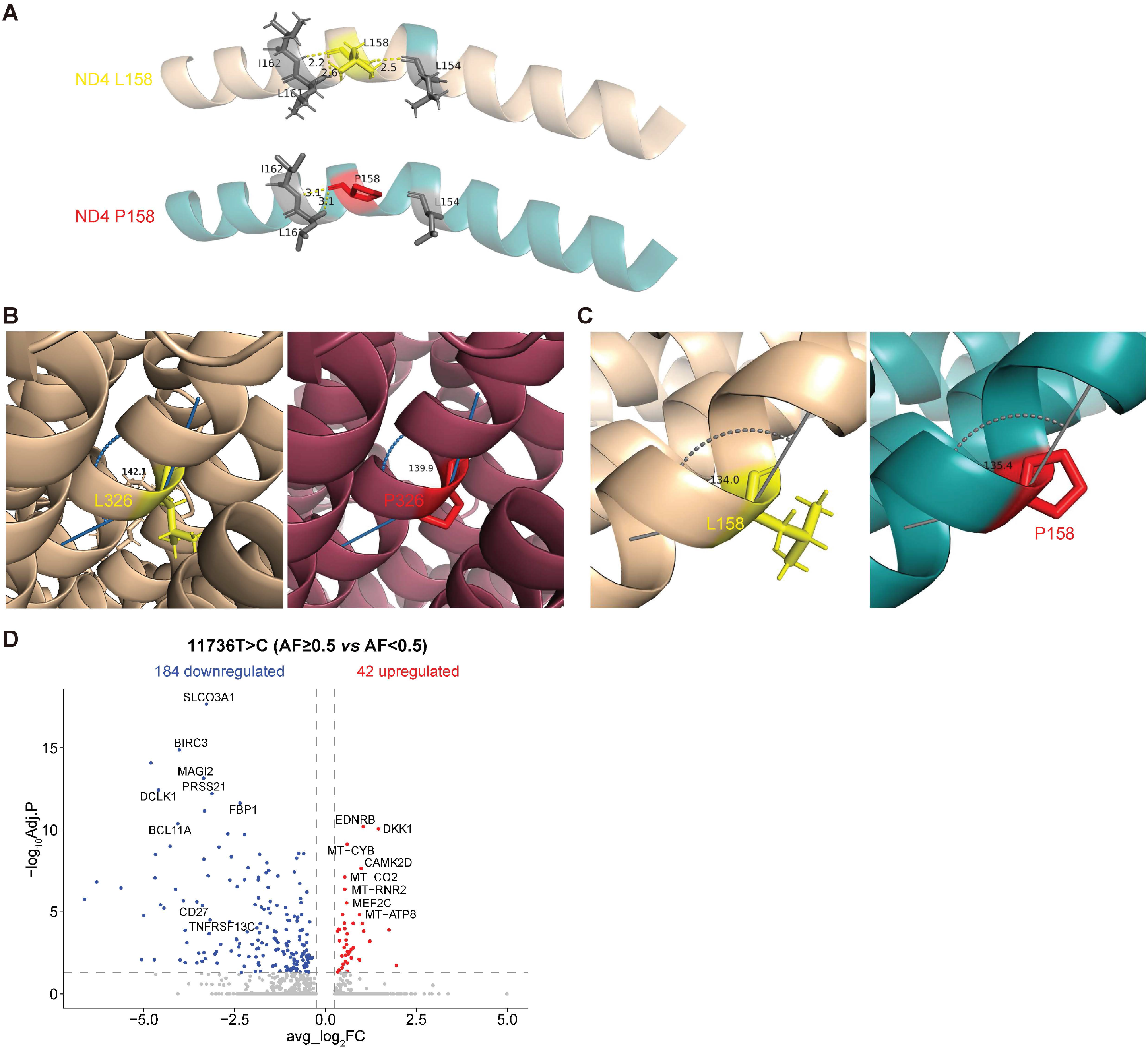
Functional and structural characterization of the 11736T>C variant (ND4, L326P). (A) Hydrogen bond interactions of L158/P158 with nearby residues in the WT (up) and mutant (bottom) ND4 helix. L158 is shown in yellow, P158 in red, and surrounding residues in grey. Hydrogen bonds are indicated by dashed lines, and residues are labeled in black. Distances (Å) between residues are also indicated. (B) Quantitative analysis of helix geometry in wild-type (WT, wheat) and mutant (L326P, raspberry) ND4. The angle (in blue) defined by the pre- and post-326 helical segments decreased from 142.1° in the WT structure to 139.9° in the mutant. (C) Quantitative analysis of helix geometry in wild-type (WT, wheat) and mutant (L158P, raspberry) ND4. The angle (in grey) defined by the pre- and post-158 helical segments increased from 134.0° in the WT structure to 135.4° in the mutant. (D) Volcano plot showing differential expressed genes between cells with high (AF≥0.5) and low (AF<0.5) AF of 11736T>C. Each dot represents a gene, with significantly upregulated genes shown in red (n=42) and downregulated genes in blue (n=184). Top upregulated and downregulated genes are labeled. Dashed lines indicate thresholds for statistical significance and fold change.

